# Development of a multi-organ single-cell atlas of systemic sclerosis reveals shared and tissue-specific pathogenic pathways

**DOI:** 10.64898/2026.10.26.761913

**Authors:** Cristian Iperi, Celina Geiss, Marija Lugar, Philip Stauffer, Nikolaos Vlachogiannis, Astrid Hofman, Alexandra Khmelevskaya, Petros P Sfikakis, Anna-Maria Hoffmann-Vold, Raphael Micheroli, Christof Seiler, Oliver Distler, Caroline Ospelt, Muriel Elhai

**Affiliations:** Center of Experimental Rheumatology, Department of Rheumatology, University Hospital Zürich, University of Zurich, Zürich, Switzerland; First Department of Propaedeutic Internal Medicine and Joint Academic Rheumatology Program, School of Medicine, National and Kapodistrian University of Athens, Athens, Greece; Department of Rheumatology, Oslo University Hospital, Oslo, Norway

**Keywords:** Systemic sclerosis, single-cell RNA sequencing, atlas, fibrosis, interferon, transcriptomics

## Abstract

**Objectives:** Systemic sclerosis (SSc) is a multi-organ autoimmune disease for which no broadly effective disease-modifying treatment exists. We developed an integrated multi-organ single-cell atlas of SSc to identify shared and organ specific disease-associated transcriptional programmes and potential therapeutic targets.

**Methods:** We integrated single-cell RNA-sequencing data from skin, lung, synovium, and peripheral blood mononuclear cells (six studies, 401,902 cells, 90 individuals) using Harmony. Pseudobulk differential expression and gene set enrichment analysis were performed at the tissue cell-compartment, cell-type, and subpopulation levels, including independent reclustering of lymphoid, myeloid and fibroblast populations.

**Results:** Disease-associated transcriptional programmes showed marked tissue dependence, even within the same cell lineage, highlighting the importance of organ-specific mechanisms in SSc. Nevertheless, recurrent extracellular matrix (ECM) remodelling programmes were identified in lung, skin and synovium, and interferon signalling programmes in skin, synovium and peripheral blood mononuclear cells, across immune and fibroblast cell populations. Although no subpopulation was exclusive to SSc, a shared myofibroblast subpopulation expressing *SFRP2, COL1A1, POSTN, ACTA2*, and *COMP* was identified across lung, skin, and synovium. In addition, perivascular and mitotic fibroblast subpopulations exhibited both profibrotic and inflammatory signatures, including interferon- and IL-2-related signalling programmes.

**Conclusions:** SSc is characterised by two complementary axes: a tissue-dependent cellular programme and a recurrent fibrotic and interferon-associated signature extending across multiple organs and cell populations. These findings are consistent with current organ-based therapeutic approaches while highlighting interferon signalling as a potential multi-organ therapeutic target. The resulting interactive single-cell atlas provides a resource for investigating shared and organ-specific disease mechanisms and guiding future therapeutic research.

Graphical abstract.
Top: study overview: 90 individuals, 6 studies and 401,902 cells across four organs: lung (7 SSc, 10 CTRL), skin (27 SSc, 10 CTRL), PBMCs (21 SSc, 6 CTRL) and synovium (6 SSc, 3 CTRL). In addition, the levels of resolution analysed (organs, cell compartments, cell types and cell subpopulations), with the number of units included at each level. Centre right: summary table of shared and organ-specific activated pathways in SSc versus CTRL across organs (plus and minus signs indicate increased and decreased activity in SSc, respectively). Centre left: pathogenic fibroblast subpopulations shared across organs (myofibroblast, perivascular and mitotic), with their activated pathways. Bottom: zoom on the cell types contributing to the shared and organ-specific programmes in each organ. SSc = systemic sclerosis; CTRL = controls; PBMCs = peripheral blood mononuclear cells; ECM = extracellular matrix/collagen metabolism; IFN = interferon signalling (IFNα-β, type I; IFNγ, type II); TNF = tumour necrosis factor signalling; IL = interleukin; ATP = ATP synthesis.

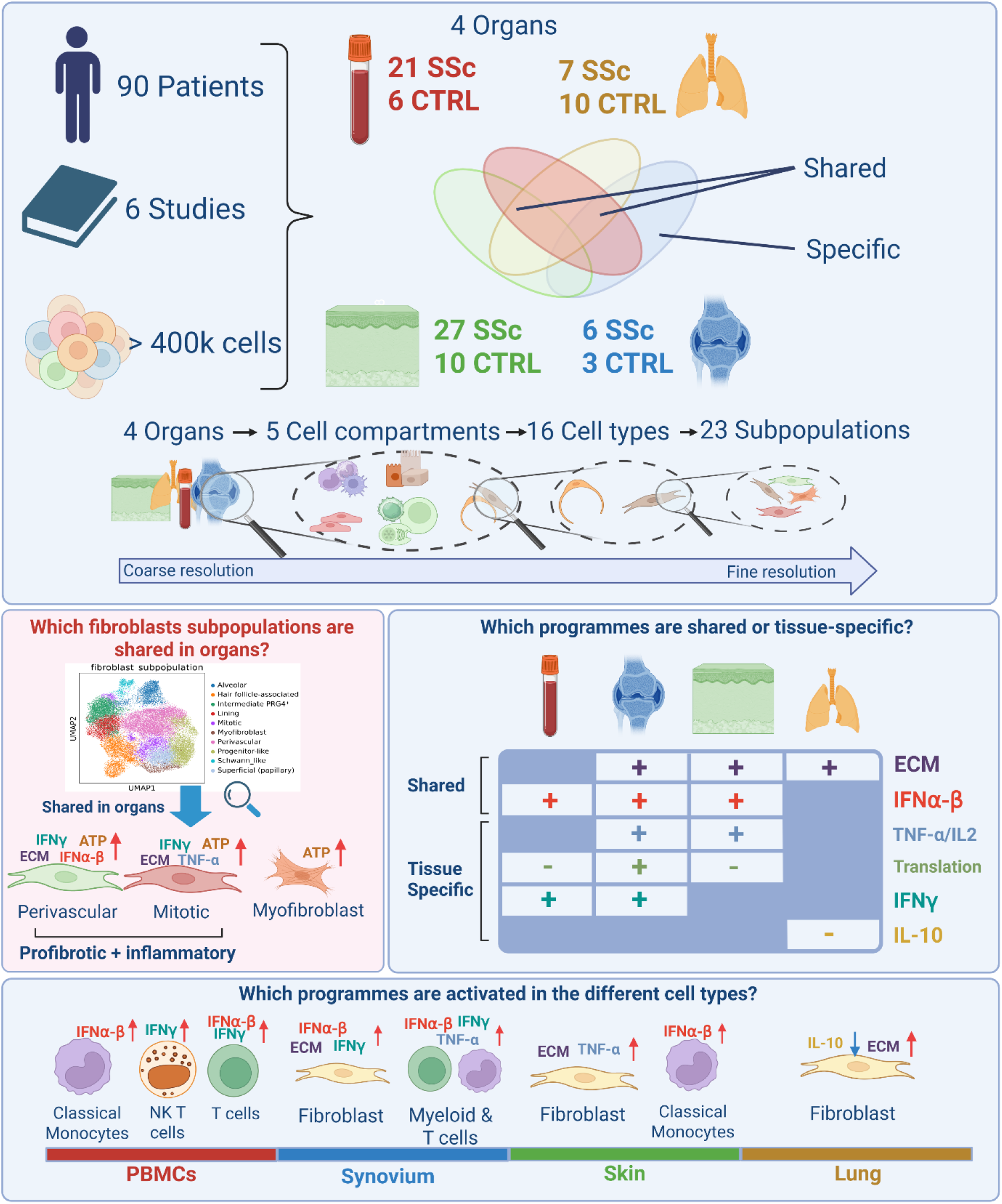

**Key messages:** *What is already known on this topic:*

❖ Single-cell studies of SSc have characterised disease-associated cell states in individual organs, but an integrated multi-organ framework identifying shared and tissue-specific mechanisms has been lacking.

*What this study adds:*

❖ This study provides the first integrated multi-organ single-cell atlas of SSc, encompassing skin, lung, synovium and peripheral blood.
❖ Disease-associated transcriptional programmes are strongly influenced by tissue context, even within the same cell lineage, highlighting the importance of organ-specific mechanisms in SSc.
❖ Beyond these tissue-specific programmes, an extracellular matrix remodelling and interferon signalling signatures were identified across multiple organs and immune and fibroblast cell populations.

*How this study might affect research, practice or policy:*

❖ These findings are consistent with the current organ-based therapeutic approach in SSc while identifying interferon signalling and extracellular matrix remodelling as potential shared therapeutic targets across multiple organs.
❖ The atlas is deployed as an open, interactive web-based platform to facilitate the exploration of shared and organ-specific disease mechanisms and guide future research.

## Introduction

Systemic sclerosis (SSc) is a prototypical autoimmune disease of inflammatory-driven fibrosis, characterised by persistent immune activation, microvascular dysfunction, and multi-organ fibrosis. Despite advances in disease management, SSc remains among the rheumatic diseases with the highest mortality and has a profound impact on quality of life, underscoring the urgent need for effective systemic therapies [1,2]. A major barrier to therapeutic progress is the limited understanding of SSc pathogenic mechanisms, further complicated by marked clinical and molecular heterogeneity among patients, a common feature of systemic autoimmune diseases [3,4]. This complexity is reflected in the limited number of effective disease-modifying therapies, with currently available treatments largely restricted to one or two specific organ manifestations [3,5–11]. Autologous haematopoietic stem cell transplantation (AHSCT) is the only intervention with demonstrated survival benefit in randomised trials, but its treatment-related morbidity and mortality restrict its use to carefully selected patients with early, rapidly progressive disease [12–14]. More recently, CAR T-cell therapies targeting autoreactive B cells have emerged as a promising therapeutic strategy for severe, refractory SSc despite limited by high cost and high morbidity [15,16].

These therapeutic limitations mirror the biological complexity of SSc. Disease progression is not driven by a single cell type, but by dysregulated interactions among immune cells, endothelial cells, and fibroblasts [17]. Understanding SSc as a systemic disease, rather than a collection of independent organ manifestations, therefore requires an analytical framework capable of simultaneously capturing cellular heterogeneity across multiple affected tissues within a unified reference.

Recent single-cell transcriptomic studies of SSc-affected tissues, including skin, lung, synovium, and peripheral blood mononuclear cells [18–24], have begun to reveal disease-associated fibroblast, endothelial, and immune cell states, as well as activation of inflammatory and fibrotic pathways. These studies have substantially advanced our understanding of tissue-level disease biology, yet they have predominantly examined individual tissues in isolation. A unified, multi-organ single-cell framework to identify shared systemic mechanisms across affected tissues is still lacking.

Therefore, we aimed to build a comprehensive, multi-organ single-cell transcriptomic atlas of SSc, integrating data from skin, lung, synovium, and peripheral blood mononuclear cell (PBMCs). We first mapped disease-associated transcriptional programmes across fibroblast, endothelial, and immune populations within each tissue. We then independently reclustered the lymphoid, myeloid and fibroblast populations across organs to determine whether conserved pathogenic cell states or molecular programmes could be identified across tissues.

## Methods

Full methodological details are provided in **Supplementary Methods**. In brief, publicly available single-cell RNA-sequencing (scRNA-seq) datasets from lung, skin, and PBMCs were retrieved from GEO/ArrayExpress (six studies in total) [18,20,21,25–27], together with an in-house synovial tissue cohort (**Table 1 and S1A**) [27]. After per-dataset quality control and preprocessing, cross-dataset comparability was assessed, and the datasets were integrated using Harmony **(Figure S1B–D)**[28].

**Table 1.** Publicly available datasets used in the present study. Datasets were retrieved from GEO/DDBJ database and integrated into the SSc atlas alongside newly generated synovium data. Abbreviations: Systemic sclerosis-associated interstitial lung disease (SSc-ILD); Diffuse cutaneous systemic sclerosis (dcSSc); limited cutaneous systemic sclerosis (lcSSc); controls (CTRL). An extended clinical table is available in Supplementary Table S1.

| **Geo/DDbj ID** | **Publication** | **Samples** | **N° Cells SSc/CTRL** | **Tissue** |
|---|---|---|---|---|
| **GSE128169** | Valenzi et al. (2019); doi: 10.1136/annrheumdis-2018-214865 | 4 SSc (SSc-ILD); 4 CTRL | 38,275; 23,712 | Lung |
| **GSE212109** | Papazoglou et al. (2022); doi: 10.1002/art.42286 | 3 SSc (SSc-ILD); 6 CTRL | 17,754; 24,250 | Lung |
| **GSE209635** | Khanna et al. (2022); doi: 10.1172/jci.insight.159566 | 15 SSc (dcSSc) | 30,628 | Skin |
| **GSE138669** | Tabib et al. (2021); doi: 10.1038/s41467-021-24607-6 | 12 SSc (dcSSc); 10 CTRL | 35,318; 26,692 | Skin |
| **E-GEAD-872** | Shimagami et al. (2025); doi: 10.1038/s41467-025-60034-7 | 21 SSc (3 dcSSc, 18 lcSSc); 6 CTRL | 135,293; 37,320 | PBMC |
| **unpublished** | Geiss, et al. (2025) doi.org/10.64898/2026.06.30.733140 [27] | 6 SSc (lcSSc); 3 CTRL | 17,857; 14,803 | Synovium |

Cell types were annotated using tissue-appropriate CellTypist [29] reference models, followed by manual curation, and grouped into five cell compartments (lymphoid, myeloid, epithelial, mesenchymal, endothelial). The integrated atlas was subsequently deployed as an interactive web-based platform comprising six analytical modules developed in Streamlit.

Differential expression analyses were performed at the patient level using a pseudobulk approach with PyDESeq2 [30], followed by gene set enrichment analysis using Reactome pathways and a Benjamini-Hochberg corrected false discovery rate (FDR) threshold of <0.05. Comparisons between SSc and controls (CTRL) were conducted at multiple levels of resolution, including organ-level pseudobulk, cell compartment, cell type and subpopulation.

Lymphoid, myeloid and fibroblast populations were extracted and independently reclustered by Leiden algorithm irrespective of organ of origin to identify disease-associated programmes shared across tissues. Low quality or mixed/doublets subclusters where then excluded. The same pseudobulk differential expression and pathway enrichment workflow was then applied within each resulting subpopulation. Finally, the pathways most consistently identified across analytical resolutions were quantified using sc.tl.score_genes. These pathways were then compared between SSc and CTRL using patient-level pseudobulk scores, Mann–Whitney U tests, and FDR.

## Results

### Construction of a multi-organ atlas for systemic sclerosis

To characterise shared and organ-specific cellular programmes in SSc, we assembled a comprehensive multi-organ scRNA-seq reference dataset integrating publicly available and in-house generated datasets spanning four sample sources: skin, lung, PBMCs, and synovium (**Table 1**; **Figure 1A and S1A**). After stringent quality control filtering, the reference included 92,638 skin cells from 37 individuals (27 dcSSc; 10 CTRL) [20,26], 103,991 lung cells from 17 individuals (7 SSc-associated interstitial lung disease; 10 CTRL) [18,19], 172,613 PBMCs from 27 individuals (21 SSc; 6 CTRL) [21], and 32,660 synovial cells from 9 individuals (6 SSc; 3 CTRL) [27], for a total of 401,902 cells (**Table 1**; **Figure 1A and S1A**). Quality control confirmed comparable sequencing depth and the absence of residual batch effects across datasets and tissues, and no additional correction was required **(Figure S1B–D)**. Cell-type annotation resolved the expected major lineages consistently across organs **(Figure 1A)**. The integrated object captured the major cellular compartments expected, including endothelial, lymphoid, epithelial, mesenchymal and myeloid compartments (**Figure 1B-C)**. Cross-tissue alignment showed mesenchymal, endothelial, and myeloid populations concordant in the UMAP embedding without discernible batch-associated segregation, supporting robust integration **(Figure S1A)**. As expected, some cell types and subpopulations were tissue specific, as monocytes in the PBMCs, synovial lining fibroblasts and skin keratinocytes. The cell compartment proportions and the demographic information of all patients and CTRL are available in **Figure S2A-B** and **Table S1**.

**Figure 1.**
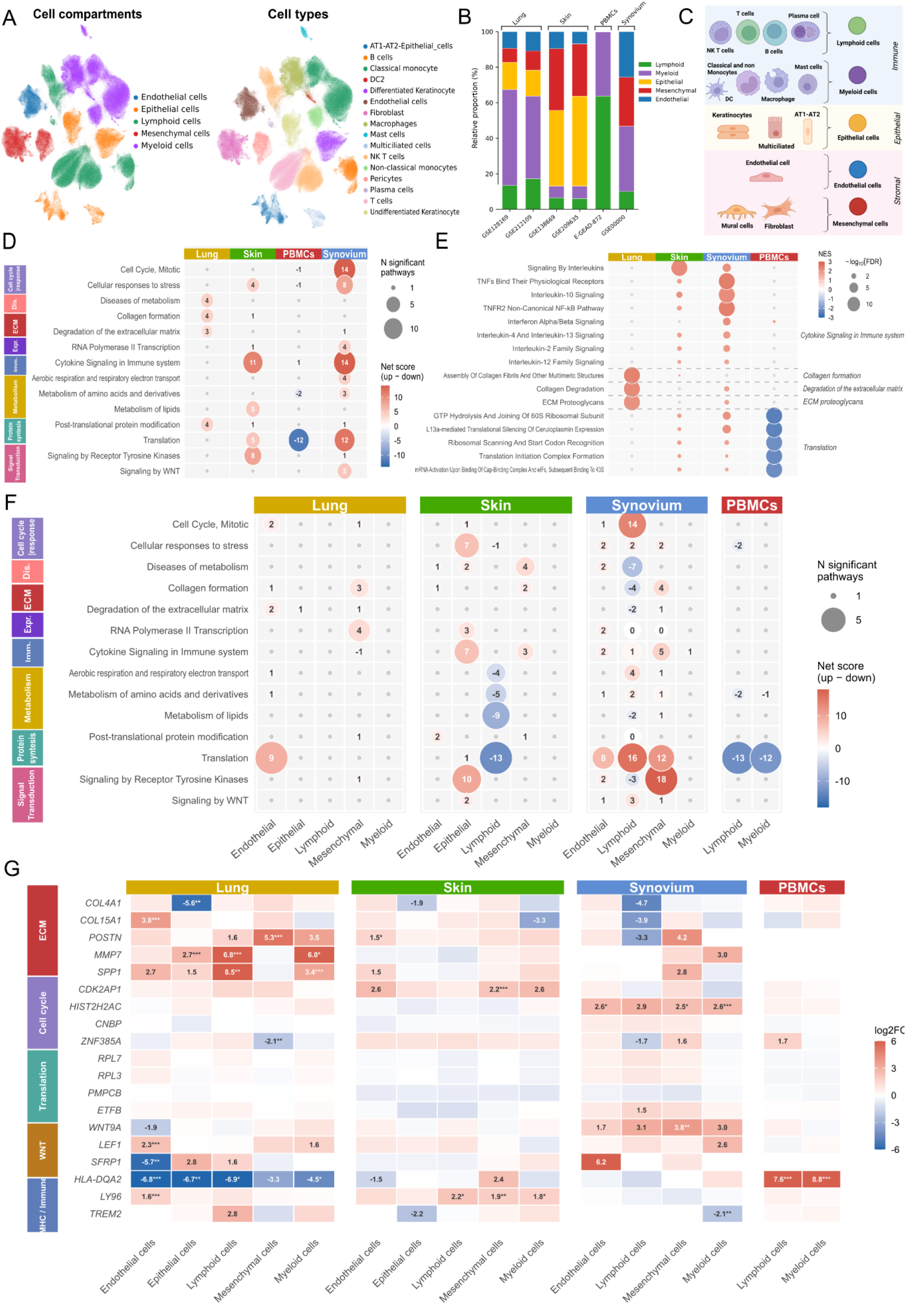
Construction and cellular composition of the multi-organ SSc single-cell atlas. **A)** UMAP (Uniform Manifold Approximation and Projection) of the integrated atlas coloured, cell compartment, and cell type. UMAP for batch and tissue in **S1A**. **B)** Relative cell compartment composition per dataset. **C)** Cell-type-to-cell-group classification: Lymphoid (T, NKT, B, plasma cells), Myeloid (monocytes, macrophages, mast cells, DC2 [type 2 dendritic cells]), Epithelial (keratinocytes, AT1-AT2 [alveolar type 1–type 2] cells, multiciliated cells), Endothelial (endothelial cells), Mesenchymal (mural cells, fibroblasts). **D)** Reactome Level 2 pathway enrichment across the four tissues pseudobulk (14 categories), grouped by parent Level 1 category (row labels; abbreviations: Cyc. = Cell cycle; Str. = Stress response; Dis. = Disease; ECM = Extracellular matrix organization; Expr = Gene expression; Imm. = Immune system; Metab. = Metabolism; Sign. = Signal transduction). Bubble size: significant pathways (FDR [false discovery rate] < 0.05); colour: net enrichment score (up − down); grey: not significant. **E)** Pathways details for Level 2 pathways in figure D related to Collagen formation/metabolism, translation and cytokine signalling. **F)** Same figure as in D showing the enriched pathways for the five cell compartments in the four-sample source. **G)** Heatmap of up to 5 genes most shared across cell compartments per pathway category (fewer where fewer than 5 met the sharing criterion), across cell compartments and tissues. Colour: log2FC (log2 fold change; SSc/CTRL); blank: not among top DE (differentially expressed) genes for that combination. Only significant genes with log2FC > 1.5 were displayed. SSc, systemic sclerosis; CTRL, control; PBMCs, peripheral blood mononuclear cells.

### Organ-level pseudobulk transcriptional signature

Having established this reference, we next asked whether the lung, skin, synovium, and blood share molecular pathways or show organ-specific activation. Gene set enrichment analysis (GSEA) was performed on pseudobulk differential expression comparing SSc to CTRL, using the Reactome gene set collection (Level 2; FDR<0.05). We first analysed all cells aggregated at the individual level within each tissue, to obtain an unbiased, organ-wide view of disease-associated pathways.

In the lung, the SSc-vs-CTRL comparison revealed a predominant enrichment of extracellular matrix (ECM) remodelling and collagen-related programmes. Similar collagen-associated signatures were observed in the skin, alongside increased tumour necrosis factor (TNF) and cytokine signalling (IL-2 and IL-20), as well as translational activity (**Figure 1D, E**). The synovium shared several of these signatures, including TNF and cytokine signalling, increased translation, and ECM remodelling, particularly ECM degradation. In addition, the synovium exhibited enrichment of both interferon type I and type II (IFN) signalling pathways. In PBMCs, IFN type I signalling was also enriched, whereas translation-related pathways were downregulated, in contrast to the increased translational activity observed in the skin and synovium. Together, these findings highlight both shared and tissue-specific transcriptional programmes in SSc. ECM and collagen-related programmes were recurrently enriched across the lung, skin, and synovium, whereas TNF/cytokine signalling and increased translational activity were shared between the skin and synovium. IFN Type I signalling emerged as a common feature of the synovium and PBMCs. Conversely, IFN type II signalling was selectively enriched in the synovium, while PBMCs displayed a distinct downregulation of translation-related pathways.

### Pathway enrichment analysis reveals SSc disease programmes within tissues

Building on this organ-level signature, we next asked which cellular compartments underlie the shared and organ-specific programmes identified above. GSEA was performed on pseudobulk differential expression comparing SSc with CTRL within each of the five cell compartments, where present, in skin, lung, synovium and PBMCs, using the Reactome gene set collection (Level 2; FDR < 0.05). The pathways most consistently shared across organs were selected (**Figure 1F**). For each of these pathways, we identified the up to five genes most consistently shared across cell compartments (**Figure 1G**).

Across the lung, skin and synovium, collagen formation emerged as one of the most consistently enriched pathways, although the magnitude of enrichment and the contributing cellular compartments differed substantially between tissues. In the lung, collagen formation was accompanied by increased cell cycle activity, with both pathways showing enrichment in endothelial and mesenchymal compartments.

Gene-level analysis further supported the enrichment of fibrotic programmes in SSc and revealed distinct cellular contributions to ECM remodelling. *POSTN* expression was increased in the mesenchymal compartment, whereas *SPP1* showed increased expression in lymphoid and myeloid compartments, potentially reflecting macrophage-associated profibrotic signalling [18] **(Figure 1G)**. Gene-level analysis further revealed distinct cellular contributions to ECM remodelling: mesenchymal cells predominantly expressed fibrillar collagens, whereas endothelial cells were enriched for basement membrane-associated collagens, including COL4A1 and COL15A1 (**Figure 1G**).

In the skin, ECM enrichment was restricted to mesenchymal and endothelial compartments, while Wnt signalling was dominant in epithelial compartment, **(Figure 1F)**, consistent with the potential role of Wnt/β-catenin signalling in keratinocytes undergoing epithelial–mesenchymal transition (EMT) in SSc skin fibrosis [31]. IFN type I signalling was also shared across epithelial and mesenchymal cell compartments, despite not being detected in the organ-level analysis.

In the synovium, collagen formation was increased in the mesenchymal compartment, while cytokine signalling predominated overall **(Figure 1F)**. Consistent with the organ pseudobulk, cytokine signalling included IFN type I signalling in mesenchymal compartment, TNF in lymphoid and myeloid compartments, and noncanonical NF-κB in lymphoid and endothelial compartments, with lymphoid compartment also showing IL-12 and IL-6. In PBMCs, the transcriptional response was more limited, dominated by downregulation of translational machinery and amino acid metabolism.

Analysis of cell compartment proportions revealed significant changes in cellular composition across SSc-affected tissues (**Figure 2A**). In the lung, the proportion of lymphoid cells was significantly reduced compared with CTRL, with trends towards expansion of the mesenchymal and epithelial compartments. In the skin, the mesenchymal compartment was significantly reduced, whereas the synovium showed trends towards a reduction in endothelial cells and an expansion of the lymphoid compartment. No significant differences were detected in PBMCs. Despite the limitation of the compositional data [32], these results suggest organ-specific alterations in cellular composition in SSc (**Figure 2B**).

**Figure 2.**
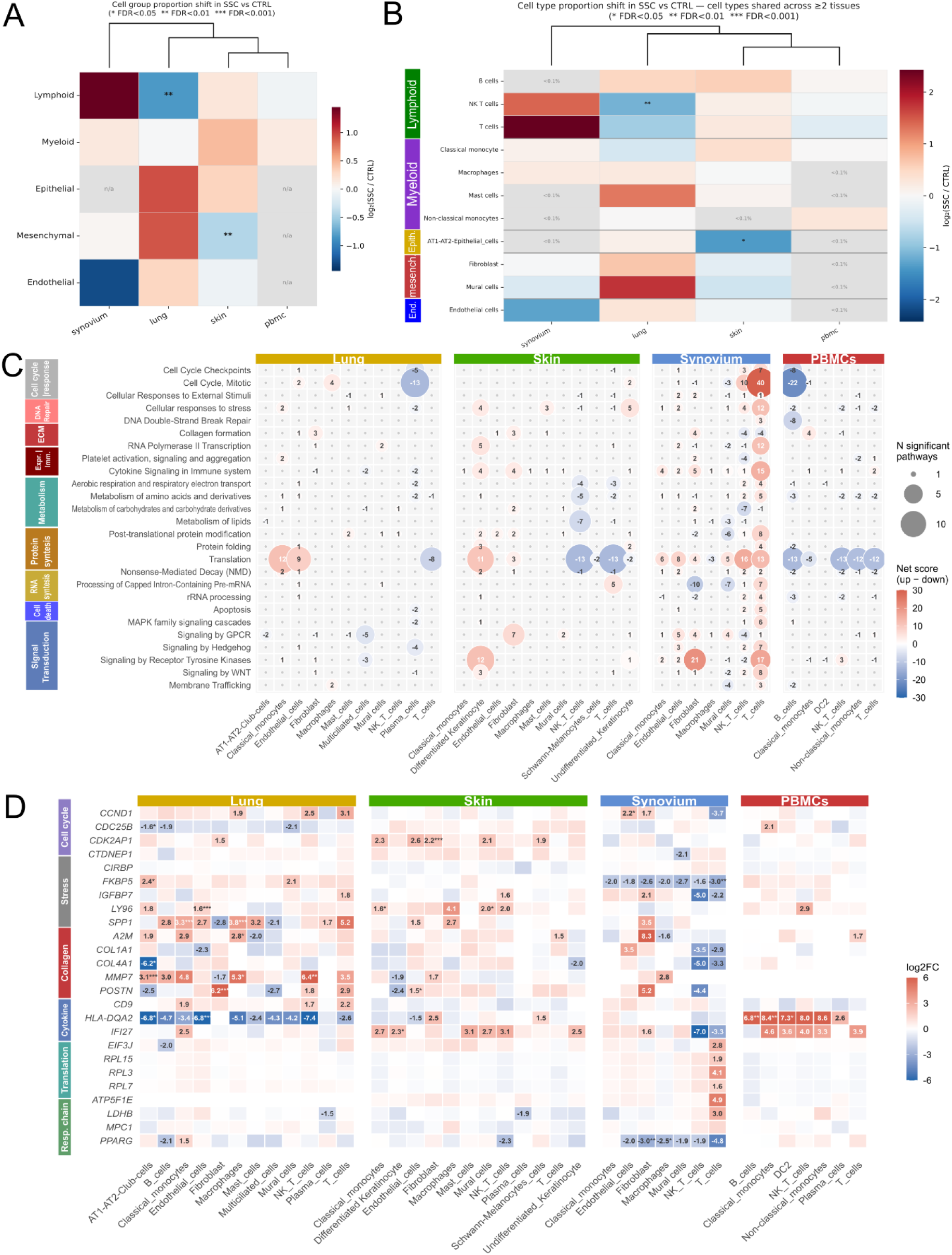
Cell-type-resolved composition and pathway landscape across SSc tissues. **A)** Cell compartments proportion shifts (log2 SSc/CTRL) across tissues, hierarchically clustered (Euclidean, average linkage). **B)** Cell type proportion shifts for types shared across ≥2 tissues, coloured by lineage (abbreviations: End. = Endothelial; Epith. = Epithelial); grey: below 0.1% abundance. Significance in A-B: Mann-Whitney U on CLR-transformed proportions, Benjamini-Hochberg corrected (* FDR<0.05, ** FDR<0.01, *** FDR<0.001). **C)** Reactome Level 2 pathway enrichment across cell types per tissue (24 categories), grouped by parent Level 1 category (row labels; abbreviations: Expr = Gene expression; Imm. = Immune system). Bubble size: significant pathways (FDR<0.05); colour: net enrichment score (up − down); displayed as a number for cells with ≥ 2 significant pathways. Grey dots: not significant. **D)** Heatmap of up to 5 genes most shared across cell types per pathway category (left strip; fewer where fewer than 5 met the sharing criterion), across cell types and tissues. Colour: log2FC (SSc/CTRL); blank: not among top DE genes for that combination. Only significant genes with log2FC > 1.5 were displayed. SSc, systemic sclerosis; CTRL, control; PBMCs, peripheral blood mononuclear cells.

At cell-type resolution, pathway enrichment analyses identified the cell types underlying these transcriptional programmes (**Figure 2C–D**). In the lung, collagen formation was enriched in endothelial cells and fibroblasts, whereas IL-10 signalling was downregulated in fibroblasts, and cell cycle pathways were enriched in macrophages. In the skin, fibroblasts showed enrichment in TNF, while IFN type I was found in keratinocytes (differentiated and undifferentiated), mast cells, classical monocytes and macrophages. In the synovium, TNF signalling was enriched in Natural Killer T (NKT) cells, T cells and macrophages, while IFN type I and II signalling was shared across T cells, pericytes, classical monocytes and fibroblasts. Cell cycle and mitotic pathways were most enriched in NKT and T cells.

In PBMCs, B cells showed the most pronounced downregulation of translation- and cell cycle-related pathways, while IFN type I signalling was specific to non-classical monocytes. At the gene level, HLA-DQA2, which encodes an MHC class II antigen-presenting molecule, was upregulated across multiple cell types in PBMC, contrasting with its downregulation in several lung cell types.

Overall, these analyses demonstrate that SSc combines recurrent molecular programmes, as cytokine signalling, translation, collagen and ECM regulation but with marked organ- and cell type-specific differences in their expression and cellular distribution.

### Tissue comparison with the SSc atlas

Having characterised the dominant programme of each organ in isolation, we next asked whether these differences primarily reflected the tissue environment or whether individual cell types retained consistent transcriptional states across organs. We therefore compared pathway enrichment and gene-level signatures for each annotated cell type across all tissues in which it was present (**Figure 3A-B**). This cross-organ comparison showed that cell identity alone was insufficient to explain the transcriptional behaviour. Several populations displayed tissue-restricted or even opposing programmes depending on their organ of origin. NKT and T cells illustrated this most starkly: amino acid metabolism was strongly suppressed in skin and PBMCs but activated to a comparable magnitude in the synovium. Translation followed a similar pattern in these cell types, being downregulated in skin, PBMCs and lung (T cells only), but not in the synovium. These findings suggest that the tissue environment could reshape lineage-specific transcriptional states. In contrast, some programmes were conserved across organs: collagen formation was concordantly upregulated in lung, synovium and skin fibroblast, as was the IFN type I signature across multiple cell types in skin, synovium and PBMCs (**Figure 3C**).

**Figure 3.**
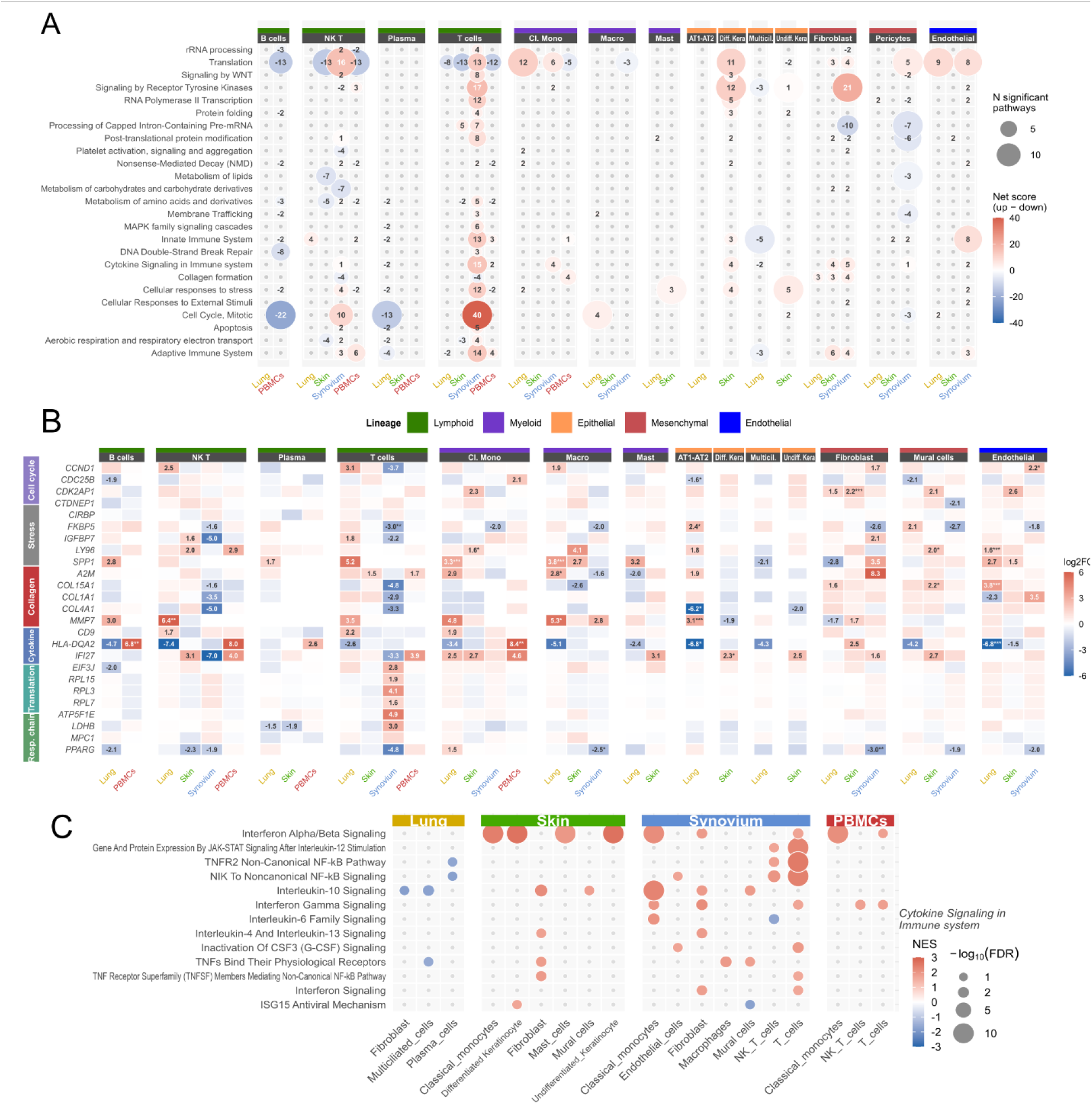
Cross-tissue pathway and gene comparison within cell types. **A)** Reactome Level 2 pathway enrichment per cell type across tissues (22 categories). Columns grouped by cell type (lineage-coloured strip) and sub-divided by tissue. Bubble size: significant pathways (FDR<0.05); colour: net enrichment score (up − down). Grey dots: not significant. **B)** Heatmap of up to 5 genes most shared across cell types per pathway category (left strip, as in Figure 1G/2D; fewer where fewer than 5 met the sharing criterion), organised by cell type (grouped by lineage) and tissue. Colour: log2FC (SSc/CTRL); blank: not among top DE genes for that combination. Only significant genes with log2FC > 1.5 were displayed. SSc, systemic sclerosis; CTRL, control; PBMCs, peripheral blood mononuclear cells. **C)** Individual Reactome pathway belonging to Cytokine Signaling in Immune system are shown for each tissue (Lung, Skin, Synovium, PBMCs), with columns representing the cell types analysed in each tissue and rows representing pathways, grouped by the Level 2 category Cytokine Signaling in Immune system (italic labels on the right, dashed lines). Only pathways significant (FDR < 0.05) in at least two cell types across the dataset (at least three for Translation) are displayed. Bubble size: −log10(FDR); colour: normalised enrichment score (NES; red = upregulated, blue = downregulated in SSc); grey dots: pathway not significant in that cell type

Overall, these results suggest that cell identity does not impose a fixed transcriptional programme; but is extensively modified by the tissue environment [33]. Given that the immune compartment showed the greatest cross-organ variability in the analyses above, and that immune compartments and fibroblast populations remain the most directly targeted by current and emerging SSc therapies [3,5,6,34–36], we focused subsequent analysis on the lymphoid, myeloid compartments and fibroblast populations. We then independently reclustered them across tissues to determine whether higher-resolution analysis could reveal conserved pathogenic subpopulations or molecular programmes that were obscured at the broader cell-type level.

### Subpopulations across the immune and fibroblast compartments

Each compartment and population was reclustered and reintegrated independently of tissue of origin, allowing subpopulations to be defined by shared transcriptional state rather than by organ.

#### Lymphoid

Reclustering of the lymphoid compartment identified six subpopulations, predominantly T and NKT cells (**Figure 4A; S3**), all shared between SSc and CTRL. PBMCs contributed most cells to every subcluster, as expected since peripheral blood is entirely immune cells, whereas lymphoid cells are only an infiltrating minority in tissues [19,20,27] (**Figure S3**). Subclustering identified naïve (MS4A1⁺, IGHD⁺) and memory (BANK1⁺, IGHD⁻) B cells, predominantly PBMC-derived, alongside CD4⁺ and CD8⁺ naïve T cells and NK/NKT cells shared between PBMCs and tissues (**Figure 4A-B; S3**). No subpopulation showed a marked SSc-vs-CTRL proportion change. GSEA on the shared PBMC/tissue subpopulations revealed increased ATP synthesis and mitochondrial cristae formation in NK and NKT cells. (**Figure 4A-B**)

**Figure 4.**
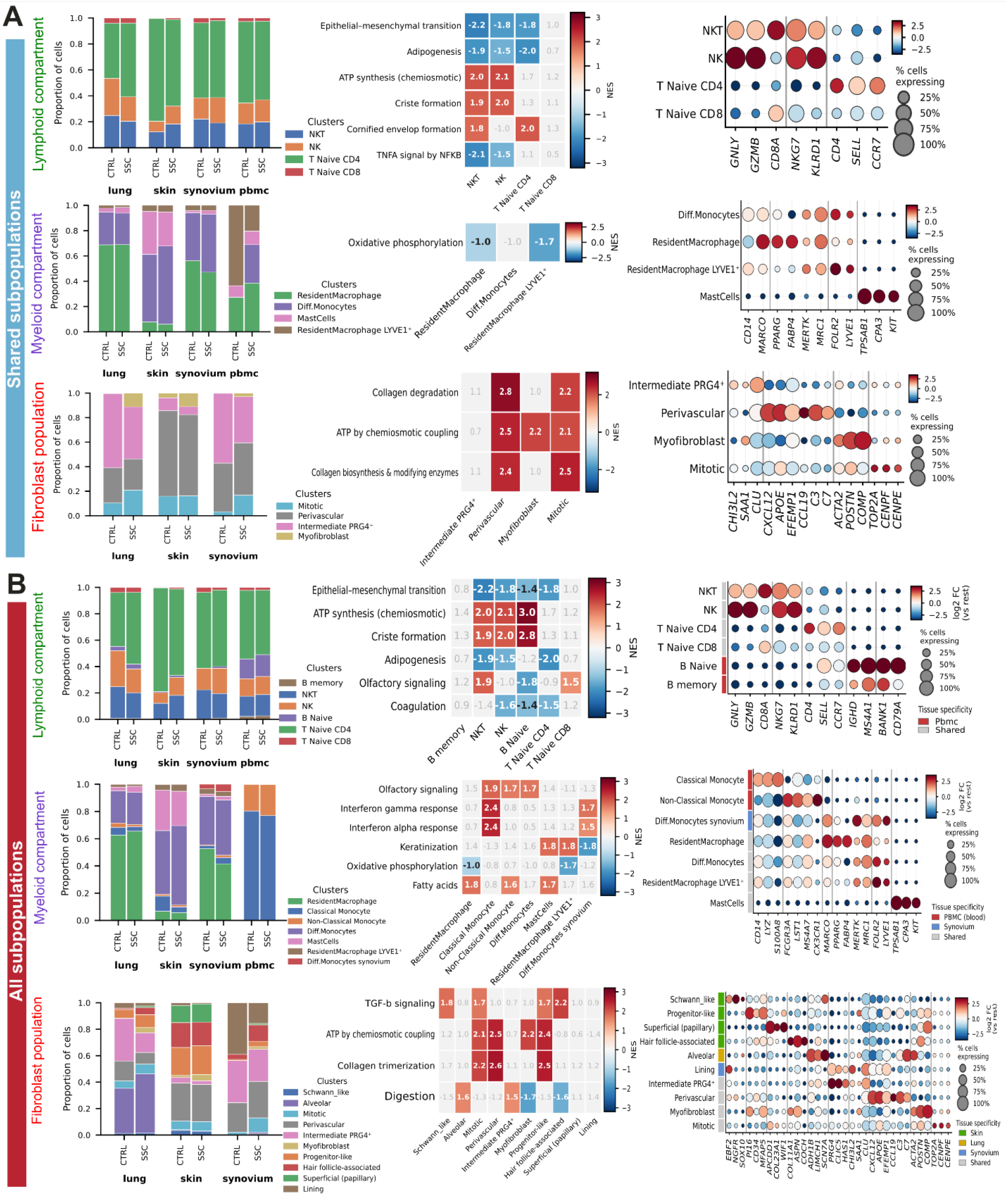
Subpopulation-level analysis of the shared and all subpopulations from lymphoid, myeloid compartments and fibroblast population. Left: Subpopulation proportions per tissue, stratified by condition: (SSc, systemic sclerosis; CTRL, control) Centre: Heatmap of the top 3 Reactome and hallmark pathways distinguishing subpopulations within each compartment and population identified from the underlying GSEA results; colour indicates the Normalised Enrichment Score (NES), with non-significant values shown in grey. Subpopulations corresponding to cell types already profiled in earlier sections were excluded from this panel to avoid redundancy. Only the pathways shared in > 1 subpopulation are shown. Right: Dotplot of marker genes characterising each subpopulation; colour indicates log2 foldchange (red: high, blue: low), dot size indicates the percentage of cells expressing the gene. **B)** Same as in A but including all subpopulations shared and tissue unique.

#### Myeloid

Reclustering of the myeloid compartment identified seven subpopulations classical monocytes, two differentiating macrophages, non-classical monocytes, mast cells, and two TREM2+ resident macrophage subpopulations, one of which, FOLR2+ LYVE1+ (**Figure 4A-B; S3**). No consistent differences in subpopulation abundance were observed between SSc and CTRL. Classical and non-classical monocytes were predominantly restricted to PBMCs, whereas differentiating macrophages were primarily associated with the synovium. The remaining subpopulations were shared across lung, skin and synovium (**Figure 4B**). Gene set enrichment analysis (GSEA) revealed reduced oxidative phosphorylation (OXPHOS) in both resident macrophage populations shared across organs, suggesting a common metabolic alteration. In contrast, interferon signalling exhibited greater tissue specificity, with IFN type I signalling enriched in classical monocytes from PBMCs and differentiating macrophages from the synovium, whereas IFN type II signalling was detected exclusively in synovial myeloid populations (**Figure 4A-B**).

#### Fibroblasts

Reclustering identified ten fibroblast subpopulations (**Figure 4A and S3**). A myofibroblast subpopulation shared across tissues was characterised by the expression of *SFRP2, COL1A1, POSTN, ACTA2* and COMP, consistent with a profibrotic phenotype. A perivascular fibroblast subpopulation expressing CXCL12, *CCL19*, *C3* and *C7* was identified suggesting an adventitial and immunoregulatory phenotype [37]. Several tissue-associated fibroblast subpopulations were identified. In the skin, these included: superficial (papillary, F1), universal (reticular, F2), hair follicle-associated (F4), progenitor-like (*MFAP5^+^; PI16^+^)* and Schwann-like fibroblasts. In the lung, an alveolar fibroblast subpopulation expressing *SCN7A* and *ADH1B* was consistent with a lung-resident fibroblast phenotype [37,38]. Synovial lining fibroblasts characterised by *PRG4* and *CLIC5* expression were exclusively detected in the synovium. In addition, two further clusters were shared across tissues: a PRG4-positive intermediate subpopulation and a mitotic subpopulation exhibiting a proliferative signature, marked by the expression of *TOP2A* and *CENPE*. A comparison between SSc and CTRL revealed a reduction in synovial lining fibroblasts and in the PRG4-positive intermediate subpopulation, whereas the lung alveolar fibroblast subpopulation was expanded in SSc (**Figure 4B**). Notably, the myofibroblast fibroblast subset expressing myofibroblast markers (*SFRP2, COL1A1, POSTN, ACTA2* and *COMP*) was expanded in SSc lung, skin and synovium. No consistent disease-associated changes in abundance were observed in the remaining subpopulations. The GSEA analysis on the shared fibroblast subpopulations revealed enrichment of collagen biosynthesis and collagen chain trimerisation pathways in perivascular and mitotic fibroblasts, together with increased ATP synthesis through chemiosmotic coupling. The myofibroblast subpopulation also exhibited increased ATP synthesis, suggesting enhanced metabolic activity in these fibroblast states (**Figure 4A-B**).

### Cross-resolution focus of fibrotic and inflammatory signatures

To complement the broader, unbiased analyses, we performed a targeted comparison of pathways that had consistently emerged as disease-associated signatures across different analytical levels. We therefore focused on key fibrotic and inflammatory pathways, including TGF-β signalling, epithelial–mesenchymal transition (EMT), collagen remodelling and selected cytokine signalling pathways (**Figure 1F, 2C**). OXPHOS was also assessed to evaluate changes in cellular energy metabolism. This was motivated by the enrichment of OXPHOS-related ATP synthesis in the mitotic, perivascular and myofibroblast populations, and by previous reports linking altered mitochondrial respiration to fibrosis in SSc fibroblasts [39]. Pathway activity scores were compared between SSc and CTRL across cell compartment, cell-type and subpopulations to identify the cellular populations contributing to these signatures and evaluate their consistency across analytical resolutions (**Figure 5A, 5B**). Collagen remodelling, EMT and TGF-β signalling were increased across most fibroblast subpopulations, with the exception of synovial lining fibroblasts. These signatures were particularly marked in the multi-organ mitotic and perivascular subpopulations (**Figure 5B**). Both subpopulations were also characterised by increased IFN type II and IL-2 signalling, with the perivascular subpopulation additionally showing IFN type I signalling, suggesting that the activation of both profibrotic and inflammatory programmes. Myofibroblasts showed a similar trend towards increased activity of fibrotic pathways, although less pronounced (**Figure 5A**). Within the myeloid compartment, IFN signalling emerged as the main cytokine-associated signature. Classical and non-classical monocytes expressed increased IFN type I and II signatures whereas monocytes predominantly showed increased IFN type II signature. These responses appeared to be mainly driven by PBMC-derived monocytes, with a potential additional contribution from differentiating monocytes in the synovium. In contrast, the lymphoid compartment was primarily characterised by increased IL-2 and IL-6 signalling within T cells, without a clearly dominant subpopulation contributing to these responses **(Figure 5A)**.

**Figure 5.**
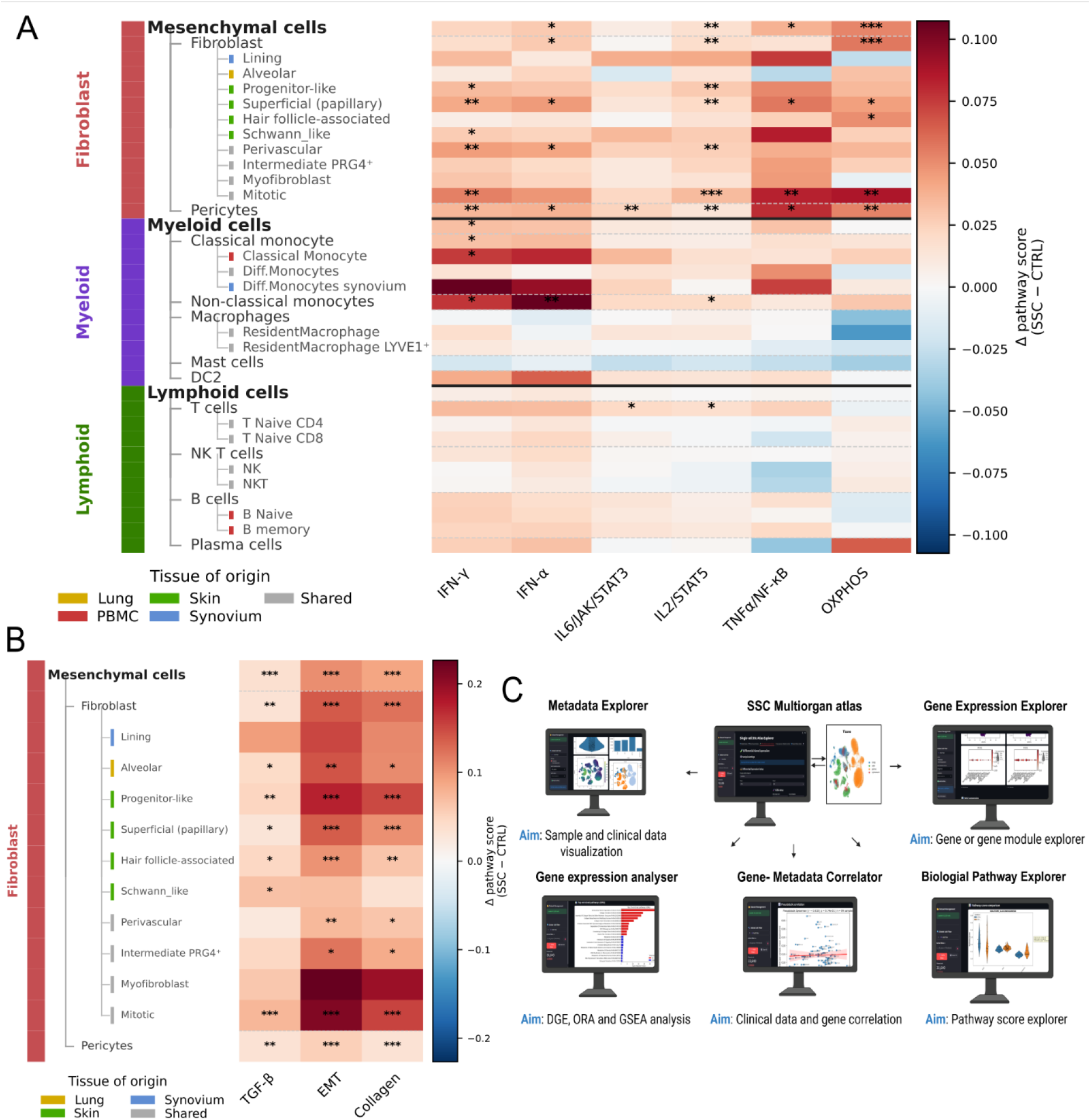
Cross-resolution pseudobulk pathway scores across cell compartments and SSc Atlas web creation. **A)** Heatmap showing the mean difference in pathway activity scores (SSc − CTRL) of lymphoid, myeloid compartments and fibroblast populations across six immune and metabolic signatures (IFN-γ, IFN-α, IL6/JAK/STAT3, IL2/STAT5, TNFα/NF-κB, and OXPHOS), stratified by cell compartment, cell type, and subpopulation. Rows are grouped hierarchically by compartment, cell type and subpopulation. Colour intensity reflects pathway score difference (red = upregulated in SSc; blue = downregulated in SSc). Significance was assessed by Wilcoxon rank-sum (Mann–Whitney) test, FDR-corrected (*p < 0.05, **p < 0.01, ***p < 0.001). The colour of each subpopulation indicates its tissue of origin (lung, yellow; skin, green; PBMCs, red; synovium, blue; shared, grey). For fibroblasts, “shared” refers to populations present in lung, skin and synovium, since PBMCs contain no fibroblasts. **B)** Heatmap showing the mean pathway score differences (SSc − CTRL) for three fibrosis-associated gene signatures: TGF-β, EMT (Epithelial-to-Mesenchymal Transition), and Collagen. Rows are ordered as in panel A, from broad cell-type categories (Fibroblast, Pericytes) to individual subpopulations. Colour scale and statistical annotation follow the same conventions as panel A. The colour of each subpopulation indicates its tissue of origin (lung, yellow; skin, green; PBMCs, red; synovium, blue; shared, grey). For fibroblasts, “shared” refers to populations present in lung, skin and synovium, since PBMCs contain no fibroblasts. **C)** Overview of the six analytical modules of the SSc Atlas web application. The atlas is accessible at [https://zisscatlas01.uzh.ch]

### Development of an interactive multi-organ SSc atlas platform

To facilitate further investigation of SSc pathogenesis, we deployed the atlas as a freely accessible, interactive web-based platform built using the Streamlit framework (**Figure 5C**). The platform enables intuitive exploration of cellular heterogeneity, gene expression, differential gene expression, pathway activity, and metadata associations across tissues and disease conditions without requiring bioinformatic expertise. Detailed descriptions of the six analytical modules provided by the platform are reported in the Methods section. The atlas is accessible at https://zisscatlas01.uzh.ch and was used as the primary analytical environment for all pathway analyses described before.

## Discussion

This study identifies two complementary axes that organise SSc pathology. The first is tissue-dependent: the same cell lineage adopts different functional programmes depending on organ context, as consistently observed across tissue, cell compartment and cell-type level of analysis. The second is transversal to cell identity: a collagen/ECM fibrotic axis in mesenchymal and endothelial compartments and an IFN/cytokine-driven axis within lymphoid, myeloid compartments and fibroblast populations across organs. Together, these findings provide a framework for understanding how pronounced tissue-specific heterogeneity coexists with shared pathogenic mechanisms in SSc.

Across levels of resolution, tissue of origin was the strongest determinant of disease-associated transcriptional programmes. The synovium was enriched for interleukin, TNF and IFN type I and II signalling across lymphoid, myeloid, mesenchymal and endothelial compartments [27]. These findings provide a biological rationale for considering TNF inhibition in SSc-associated arthritis, although its use remains controversial because of concerns regarding potential worsening of pulmonary fibrosis [40]. Furthermore, Wnt/β-catenin signalling was detected in lymphoid, endothelial and mesenchymal compartments in the synovium. This is compatible with the established role of WNT signalling in fibroblast activation, myofibroblast differentiation, and extracellular matrix production in SSc [41].

The skin exhibited a partially overlapping inflammatory profile, with IFN type I signalling identified in both epithelial and mesenchymal compartments, whereas TNF signature was predominantly associated to fibroblasts and Wnt/β-catenin signalling with epithelial compartment. In contrast, PBMCs showed a distinct transcriptional profile characterised by downregulation of the translational machinery as observed in other autoimmune diseases [42]. The lung showed a more restricted cytokine signature, characterised by reduced IL-10 signature. Importantly, even within the same cell lineage, disease-associated transcriptional programmes varied considerably across organs. For instance, NKT and T cells exhibited downregulation of amino acid metabolism pathways in the skin and peripheral blood, but upregulation of these pathways in the synovium. This tissue-dependent heterogeneity was also identified at the individual gene level. A representative example is HLA-DQA2, gene involved in antigen presentation whose locus has been genetically associated with SSc susceptibility [43], While *HLA-DQA2* was downregulated across pulmonary cell compartments, it was among the most upregulated genes in PBMCs. Together, these findings highlight the pronounced influence of tissue context on disease-associated transcriptional programmes, consistent with current organ-based therapeutic approaches in SSc [3].

However, this tissue dominance was not absolute. Cross-resolution analyses identified two programmes recurred across resolution levels in several organs as previously mentioned. The first was a fibroblast collagen/ECM fibrotic programme. In the lung, skin and synovium, ECM organisation and collagen-related pathways dominated, consistent with the literature [24], with a division of labour in collagen metabolism: fibroblasts primarily expressed fibrillar collagen and endothelial cells (in skin and lung) contributed basement membrane-associated components.

The second was IFN signalling, where IFN type I was detected in skin, synovium and blood across multiple resolution levels, extending previous independent analysis on IFN signature in SSc synovitis [27]. This supports IFN signalling as a cross-organ therapeutic target, consistent with the ongoing anifrolumab trials in SSc [20,44]. More broadly, a shared immune-fibroblast axis provides a biological rationale for disease-modifying strategies with multi-organ efficacy, such as haematopoietic stem cell transplantation and CD19-targeted CAR T cell therapy [12–15]. Independent reclustering of the lymphoid, myeloid and fibroblast populations, performed irrespective of tissue origin, enabled the identification of distinct cellular subpopulations contributing to these shared disease-associated programmes. IFN type I and II signalling was most pronounced in a classical monocyte subpopulation predominantly derived from PBMCs, whereas multi-organ resident macrophage subpopulations exhibited reduced OXPHOS, suggesting that IFN activation and metabolic alterations preferentially affect distinct myeloid subset. Within the fibroblast population, collagen/ECM remodelling signalling were particularly enriched in multi-organ perivascular and mitotic subpopulations, together with increased OXPHOS-related ATP generation. This is consistent with experimental studies showing that respiratory-chain inhibition reduces collagen secretion in dermal fibroblasts [39,47,48]. Targeting OXPHOS in these subpopulations may therefore warrant investigation, given the antifibrotic effect of mitochondrial inhibition in experimental SSc models [49,50].

Despite the identification of shared disease-associated programmes, none of the reclustered subpopulations was exclusive to SSc or to CTRL. Instead, differences occurred within pre-existing subpopulations rather than through the appearance of entirely new disease-specific cell states. The fibroblast population represented a partial exception: the myofibroblast subcluster was consistently expanded in all organs in SSc and, together with the perivascular and mitotic subpopulations, showed enrichment of OXPHOS-related ATP synthesis. The perivascular and mitotic subpopulations additionally showed the strongest and most coherent disease-associated pathway profile (ECM remodelling and inflammation signalling), making these three subpopulations the closest candidates to pathogenic states in this atlas. Overall, these findings suggest that SSc pathogenesis may be driven at least in part, by the dysregulation of functional programmes within shared immune and fibroblast population rather than by the emergence of disease-specific cell states. Several limitations should be acknowledged. The atlas integrates publicly available datasets with heterogeneous protocols and sequencing depths, which may introduce residual technical variability despite batch correction. Furthermore, the pronounced collagen/extracellular matrix signature in the lung may partly reflect disease stage rather than an organ-intrinsic property, as lung samples were end-stage explants whereas synovium, PBMC and skin cohorts were largely early-stage. Third, the relatively small number of synovial CTRL limited the statistical power of compositional analyses. Finally, the observational nature of single-cell transcriptomic analyses does not allow causal relationships to be established, and the identified pathways require further functional validation.

However, this study has several strengths: this work establishes, to our knowledge, the first integrated multi-organ single-cell atlas of SSc integrating skin, lung, synovium, and blood within a unified reference. This integration within a unified analytical framework enabled systematic comparisons of disease-associated transcriptional programmes across organs and cellular populations. Moreover, the multi-resolution approach, extending from tissue-level analyses to individual cell types and subpopulations, allowed the identification of both tissue-specific and recurrent molecular signatures that might otherwise remain undetected in single-organ studies. The development of an interactive atlas further provides a valuable resource for the research community, facilitating data exploration and hypothesis generation.

## Conclusion

In conclusion, this study highlights the coexistence of pronounced tissue-dependent transcriptional heterogeneity and recurrent molecular programmes in SSc, including collagen/extracellular matrix remodelling and IFN signalling across multiple organs and immune and fibroblast populations. These findings provide a framework for understanding shared and organ-specific disease mechanisms and highlight potential therapeutic pathways for future investigation. The resulting interactive single-cell atlas represents a resource to support further research into SSc pathogenesis and the identification of potential therapeutic targets.

## Supporting information

Supplementary Material. Includes the supplementary tables and figures.

## Ethics approval

This study involves only publicly available data. All publicly available datasets used in this study (skin, lung, PBMCs, synovium) were generated under the ethical approvals reported in their original publications.

## Funding sources

This work was supported by the UZH Advanced Clinician Scientist Program, awarded to ME which covered the salary of CI. It was further supported by a Graduate Campus (GRC) Short Grant from the University of Zurich (Grant No. SG_323), awarded to CI, which supported the development and online deployment of the interactive atlas platform.

## Data availability statement

Publicly available datasets were retrieved from the Genomic Expression Archive (GEA/DDBJ) and the Gene Expression Omnibus (GEO). Lung: GSE128169 (Valenzi et al., 2019) and GSE212109 (Papazoglou et al., 2022). Skin: GSE209635 (Khanna et al., 2022) and GSE138669 (Tabib et al., 2021). PBMCs: E-GEAD-872 (Shimagami et al., 2025). Synovial tissue data are not yet publicly deposited; they will be made available upon publication of the originating manuscript (Geiss et al., 2026 [27]). Code used for data integration, analysis, and generation of the interactive atlas platform is available at: https://github.com/IxI-97/Atlas_SSC.

## Declaration of competing interests

C. Iperi: none declared. C. Geiss: none declared. M. Lugar: none declared. P. Stauffer: none declared. N. Vlachogiannis: none declared. A. Hofman: none declared. R. Micheroli: received speaker fees from AbbVie, Johnson & Johnson, UCB, and Eli LillyA. Khmelevskaya: none declared. P.P. Sfikakis: reports grants and personal fees from Abbvie, Boehringer Ingelheim, Eli-Lilly, MSD, Pfizer and UCB outside the submitted work. A-M. Hoffmann-Vold: consultancy/speaker fees from Boehringer Ingelheim, Janssen, Medscape, Merck Sharp & Dohme, Novartis, Roche, AbbVie, Avalyn, AstraZeneca, Bristol Myers Squibb, Calluna Pharma, Genentech, Pliant, Werfen, Oliver Distler, Kymera, Mitsubishi Tanabe and UCB. C. Seiler: none declared. O. Distler: consultancy/speaker fees from Boehringer Ingelheim, 4P-Pharma, AbbVie, Acepodia, Aera, AnaMar, Anaveon, Argenx, AstraZeneca, Avalyn, BMS, Calluna, Cantargia, CSL Behring, EMD Serono, Galderma, Galapagos, Gossamer, Hemetron, Innovaderm, Kali, Lilly, Mediar, MSD Merck, Nkarta, Novartis, Oorja Bio, Orion, Pliant, Prometheus, Quell, Scleroderma Research Foundation, Skyhawk, Tandem, Topadur, UCB and Umlaut.bio. C. Ospelt: none declared. M. Elhai: grant/research support from Vontobel Stiftung, Pfizer, Novartis Foundation for Bio-Medical Research, Iten Kohaut Foundation, Kurt und Senta Herrmann Foundation, Foundation for Research in Rheumatology (FOREUM), and University of Zurich; congress support from AstraZeneca; speaker and consultancy fees from Boehringer Ingelheim.

## Patient and public involvement

The project was presented and discussed with A. Eisenring and J. Messmer, two patient research partners involved in the synovitis project [27] during one of the regular consultation meetings, which are also used to identify research questions and topics of priority to the SSc patient community. This study addresses one such question, previously raised by our patient research partners concerning multi-organ involvement and development of disease modifying approach.

## Provenance and peer review

Not commissioned, externally peer reviewed.

## Declaration of generative AI use

Generative AI (Claude, Anthropic) was used to assist with language editing and proofreading of the manuscript text, and to assist with the development and modification of R and python scripts used for figure generation. All AI-assisted output was reviewed, verified, and edited by the authors, who take full responsibility for the accuracy and integrity of the final manuscript and code.

## Contributorship (CRediT)

Conceptualization: M. Elhai. Methodology: C. Iperi, C. Geiss. Software: C. Iperi, C. Geiss. Validation: C. Iperi, A. Khmelevskaya, A. Hofman, N. Vlachogiannis, M. Elhai, O. Distler, C. Ospelt, P.P. Sfikakis, C. Seiler. Formal analysis: C. Iperi. Investigation: C. Geiss. Resources: M. Elhai, O. Distler, C. Ospelt, M. Lugar. Data curation: C. Iperi, C. Geiss, C. Seiler. Writing – original draft: C. Iperi. Writing – review & editing: all authors. Visualization: C. Iperi, M. Elhai, C. Geiss, C. Ospelt, A. Hofman, A. Khmelevskaya, M. Lugar, P. Stauffer. Supervision: M. Elhai. Project administration: M. Elhai, C. Ospelt. Funding acquisition: M. Elhai, C. Iperi.

## Acknowledgements

The authors thank the IT department of the University of Zurich (UZH) for their assistance in deploying the interactive atlas online. We thank our patient research partners A. Eisenring and J. Messmer for ensuring the study is targeting patients’needs.

## Notes

https://zisscatlas01.uzh.ch

## References

1 Good SD, Lee JY, Johnson RE, et al. A scoping review of the epidemiology of systemic sclerosis and its organ manifestations: 2018-2024. Curr Opin Rheumatol. 2025;37:103–12. doi: 10.1097/BOR.0000000000001063

2 Elhai M, Meune C, Boubaya M, et al. Mapping and predicting mortality from systemic sclerosis. Ann Rheum Dis. 2017;76:1897–905. doi: 10.1136/annrheumdis-2017-211448

3 Del Galdo F, Lescoat A, Conaghan PG, et al. EULAR recommendations for the treatment of systemic sclerosis: 2023 update. Ann Rheum Dis. 2025;84:29–40. doi: 10.1136/ard-2024-226430

4 Barturen G, Babaei S, Català-Moll F, et al. Integrative Analysis Reveals a Molecular Stratification of Systemic Autoimmune Diseases. Arthritis Rheumatol. 2021;73:1073–85. doi: 10.1002/art.41610

5 Khanna D, Lin CJF, Furst DE, et al. Tocilizumab in systemic sclerosis: a randomised, double-blind, placebo-controlled, phase 3 trial. Lancet Respir Med. 2020;8:963–74. doi: 10.1016/S2213-2600(20)30318-0

6 Tashkin DP, Roth MD, Clements PJ, et al. Mycophenolate mofetil versus oral cyclophosphamide in scleroderma-related interstitial lung disease (SLS II): a randomised controlled, double-blind, parallel group trial. Lancet Respir Med. 2016;4:708–19. doi: 10.1016/S2213-2600(16)30152-7

7 Zhu JL, Black SM, Chen HW, et al. Emerging treatments for scleroderma/systemic sclerosis. Fac Rev. 2021;10:43. doi: 10.12703/r/10-43

8 Distler O, Highland KB, Gahlemann M, et al. Nintedanib for Systemic Sclerosis-Associated Interstitial Lung Disease. N Engl J Med. 2019;380:2518–28. doi: 10.1056/NEJMoa1903076

9 Ebata S, Yoshizaki A, Oba K, et al. Safety and efficacy of rituximab in systemic sclerosis (DESIRES): open-label extension of a double-blind, investigators-initiated, randomised, placebo-controlled trial. Lancet Rheumatol. 2022;4:e546–55. doi: 10.1016/S2665-9913(22)00131-X

10 Khanna D, Spino C, Johnson S, et al. Abatacept in Early Diffuse Cutaneous Systemic Sclerosis: Results of a Phase II Investigator-Initiated, Multicenter, Double-Blind, Randomized, Placebo-Controlled Trial. Arthritis Rheumatol. 2020;72:125–36. doi: 10.1002/art.41055

11 Khanna D, Lin CJF, Furst DE, et al. Tocilizumab in systemic sclerosis: a randomised, double-blind, placebo-controlled, phase 3 trial. Lancet Respir Med. 2020;8:963–74. doi: 10.1016/S2213-2600(20)30318-0

12 Sieiro Santos C. Autologous Hematopoietic Stem Cell Transplantation in Systemic Sclerosis: Current Evidence and Future Directions. ACR Open Rheumatol. 2025;7:e70131. doi: 10.1002/acr2.70131

13 van Laar JM, Farge D, Sont JK, et al. Autologous hematopoietic stem cell transplantation vs intravenous pulse cyclophosphamide in diffuse cutaneous systemic sclerosis: a randomized clinical trial. JAMA. 2014;311:2490–8. doi: 10.1001/jama.2014.6368

14 Burt RK, Shah SJ, Dill K, et al. Autologous non-myeloablative haemopoietic stem-cell transplantation compared with pulse cyclophosphamide once per month for systemic sclerosis (ASSIST): an open-label, randomised phase 2 trial. Lancet. 2011;378:498–506. doi: 10.1016/S0140-6736(11)60982-3

15 Auth J, Müller F, Völkl S, et al. CD19-targeting CAR T-cell therapy in patients with diffuse systemic sclerosis: a case series. Lancet Rheumatol. 2025;7:e83–93. doi: 10.1016/S2665-9913(24)00282-0

16 Bergmann C, Müller F, Distler JHW, et al. Treatment of a patient with severe systemic sclerosis (SSc) using CD19-targeted CAR T cells. Ann Rheum Dis. 2023;82:1117–20. doi: 10.1136/ard-2023-223952

17 Truchetet ME, Brembilla NC, Chizzolini C. Current Concepts on the Pathogenesis of Systemic Sclerosis. Clin Rev Allergy Immunol. 2023;64:262–83. doi: 10.1007/s12016-021-08889-8

18 Papazoglou A, Huang M, Bulik M, et al. Epigenetic Regulation of Profibrotic Macrophages in Systemic Sclerosis-Associated Interstitial Lung Disease. Arthritis Rheumatol. 2022;74:2003–14. doi: 10.1002/art.42286

19 Valenzi E, Tabib T, Papazoglou A, et al. Disparate Interferon Signaling and Shared Aberrant Basaloid Cells in Single-Cell Profiling of Idiopathic Pulmonary Fibrosis and Systemic Sclerosis-Associated Interstitial Lung Disease. Front Immunol. 2021;12:595811. doi: 10.3389/fimmu.2021.595811

20 Khanna D, Padilla C, Tsoi LC, et al. Tofacitinib blocks IFN-regulated biomarker genes in skin fibroblasts and keratinocytes in a systemic sclerosis trial. JCI Insight. 2022;7:e159566. doi: 10.1172/jci.insight.159566

21 Shimagami H, Nishimura K, Matsushita H, et al. Single-cell analysis reveals immune cell abnormalities underlying the clinical heterogeneity of patients with systemic sclerosis. Nat Commun. 2025;16:4949. doi: 10.1038/s41467-025-60034-7

22 Gur C, Wang S-Y, Sheban F, et al. LGR5 expressing skin fibroblasts define a major cellular hub perturbed in scleroderma. Cell. 2022;185:1373–1388.e20. doi: 10.1016/j.cell.2022.03.011

23 Rius Rigau A, Xu M, Liu Z, et al. Deep phenotyping of skin tissue remodeling in patients with systemic sclerosis treated with CD19-CAR T cells. Nat Commun. 2026;17:4640. doi: 10.1038/s41467-026-72817-7

24 Zielonka J, Li N, Liu Y, et al. Skin Transcriptomics Reveal Shared Molecular Mechanisms for Skin and Lung Involvement in Systemic Sclerosis. Arthritis Rheumatol. Published Online First: 3 April 2026. doi: 10.1002/art.70163

25 Valenzi E, Bulik M, Tabib T, et al. Single-cell analysis reveals fibroblast heterogeneity and myofibroblasts in systemic sclerosis-associated interstitial lung disease. Ann Rheum Dis. 2019;78:1379–87. doi: 10.1136/annrheumdis-2018-214865

26 Tabib T, Huang M, Morse N, et al. Myofibroblast transcriptome indicates SFRP2hi fibroblast progenitors in systemic sclerosis skin. Nat Commun. 2021;12:4384. doi: 10.1038/s41467-021-24607-6

27 Geiss C, Cebrian CC, Houtman M, et al. Synovitis in systemic sclerosis is an interferon-driven stromal condition distinct from rheumatoid arthritis. 2026;2026.06.30.733140.

28 Korsunsky I, Millard N, Fan J, et al. Fast, sensitive and accurate integration of single-cell data with Harmony. Nat Methods. 2019;16:1289–96. doi: 10.1038/s41592-019-0619-0

29 Domínguez Conde C, Xu C, Jarvis LB, et al. Cross-tissue immune cell analysis reveals tissue-specific features in humans. Science. 2022;376:eabl5197. doi: 10.1126/science.abl5197

30 Muzellec B, Teleńczuk M, Cabeli V, et al. PyDESeq2: a python package for bulk RNA-seq differential expression analysis. Bioinformatics. 2023;39:btad547. doi: 10.1093/bioinformatics/btad547

31 Nagib L, Kumar AS, Stratton R. Understanding the Role of Epithelial Cells in the Pathogenesis of Systemic Sclerosis. Cells. 2025;14:962. doi: 10.3390/cells14130962

32 Büttner M, Ostner J, Müller CL, et al. scCODA is a Bayesian model for compositional single-cell data analysis. Nat Commun. 2021;12:6876. doi: 10.1038/s41467-021-27150-6

33 Domínguez Conde C, Xu C, Jarvis LB, et al. Cross-tissue immune cell analysis reveals tissue-specific features in humans. Science. 2022;376:eabl5197. doi: 10.1126/science.abl5197

34 Distler JHW, Kuwana M, Assassi S, et al. Emerging therapies for the treatment of systemic sclerosis. Nat Rev Rheumatol. 2025;21:612–25. doi: 10.1038/s41584-025-01294-x

35 Mulcaire-Jones E, Low AHL, Domsic R, et al. Advances in biological and targeted therapies for systemic sclerosis. Expert Opin Biol Ther. 2023;23:325–39. doi: 10.1080/14712598.2023.2196009

36 Naveed E, Lescoat A, Matucci-Cerinic M, et al. Emerging drugs in phase 2/3 and selected phase 1/2 cellular-therapy trials: a glimpse into the future of systemic sclerosis treatment. Ann Rheum Dis. 2026;S0003–4967(26)00593-5. doi: 10.1016/j.ard.2026.09.015

37 Steele L, Olabi B, Roberts K, et al. A single-cell and spatial genomics atlas of human skin fibroblasts reveals shared disease-related fibroblast subtypes across tissues. Nat Immunol. 2025;26:1807–20. doi: 10.1038/s41590-025-02267-8

38 Travaglini KJ, Nabhan AN, Penland L, et al. A molecular cell atlas of the human lung from single-cell RNA sequencing. Nature. 2020;587:619–25. doi: 10.1038/s41586-020-2922-4

39 Cantanhede IG, Liu H, Balbuena Rodriguez V, et al. Exploring metabolism in scleroderma reveals opportunities for pharmacological intervention for therapy in fibrosis. Front Immunol. 2022;13. doi: 10.3389/fimmu.2022.1004949

40 Distler JHW, Jordan S, Airo P, et al. Is there a role for TNFα antagonists in the treatment of SSc? EUSTAR expert consensus development using the Delphi technique. Clin Exp Rheumatol. 2011;29:S40–45.

41 Wei J, Fang F, Lam AP, et al. Wnt/β-catenin signaling is hyperactivated in systemic sclerosis and induces Smad-dependent fibrotic responses in mesenchymal cells. Arthritis Rheum. 2012;64:2734–45. doi: 10.1002/art.34424

42 Iperi C, Rekha AA, Arvidsson G, et al. Interferon-α-Driven Stratification of B Cells Reveals Metabolic Reprogramming of Double-Negative, Naïve, and Transitional Cell Subsets and Refines Molecular Classification in Sjögren’s Disease. Eur J Immunol. 2026;56:e70275. doi: 10.1002/eji.70275

43 Acosta-Herrera M, Kerick M, Lopéz-Isac E, et al. Comprehensive analysis of the major histocompatibility complex in systemic sclerosis identifies differential HLA associations by clinical and serological subtypes. Ann Rheum Dis. 2021;80:1040–7. doi: 10.1136/annrheumdis-2021-219884

44 Radić M, Prižmić PŠ, Bečić T, et al. Anifrolumab—A Potential New Systemic Sclerosis Treatment. J Clin Med. 2026;15:1104. doi: 10.3390/jcm15031104

45 Nevskaya T, Zheng B, Baxter CA, et al. Skin improvement is a surrogate for favourable changes in other organ systems in early diffuse cutaneous systemic sclerosis. Rheumatology (Oxford). 2020;59:1715–24. doi: 10.1093/rheumatology/kez529

46 Zheng B, Nevskaya T, Baxter CA, et al. Changes in skin score in early diffuse cutaneous systemic sclerosis are associated with changes in global disease severity. Rheumatology (Oxford). 2020;59:398–406. doi: 10.1093/rheumatology/kez299

47 Darawsha A, Trachtenberg A, Sharoni Y. ARE/Nrf2 Transcription System Involved in Carotenoid, Polyphenol, and Estradiol Protection from Rotenone-Induced Mitochondrial Oxidative Stress in Dermal Fibroblasts. Antioxidants (Basel). 2024;13:1019. doi: 10.3390/antiox13081019

48 Zhou X, Trinh-Minh T, Tran-Manh C, et al. Impaired Mitochondrial Transcription Factor A Expression Promotes Mitochondrial Damage to Drive Fibroblast Activation and Fibrosis in Systemic Sclerosis. Arthritis & Rheumatology. 2022;74:871–81. doi: 10.1002/art.42033

49 Ursini F, Grembiale RD, D’Antona L, et al. Oral Metformin Ameliorates Bleomycin-Induced Skin Fibrosis. J Invest Dermatol. 2016;136:1892–4. doi: 10.1016/j.jid.2016.05.097

50 Karatas A, Oz B, Celik C, et al. Tofacitinib and metformin reduce the dermal thickness and fibrosis in mouse model of systemic sclerosis. Sci Rep. 2022;12:2553. doi: 10.1038/s41598-022-06581-1

