## Supplementary Material. Includes the supplementary tables and figures. for "Development of a multi-organ single-cell atlas of systemic sclerosis reveals shared and tissue-specific pathogenic pathways"

**Supplementary Table S1. Clinical characteristics of SSc patients across integrated datasets**

| Tissue | Dataset | N (SSc) | Age (yrs) | Female (%) | Subtype | Disease duration (yrs) | mRSS | FVC%/DLCO% predicted | Autoantibodies | Organ involvement | Immunosuppression at sampling | Population context |
| --- | --- | --- | --- | --- | --- | --- | --- | --- | --- | --- | --- | --- |
| Lung | Valenzi et al. 2019 (GSE128169) | 4 (8 samples, upper+lower lobe) | 56.8 [43–67] | 25.0 | SSc-ILD 100% | NR | NR | NR | NR | All UIP with acute lung injury / lymphoid aggregates; mPAP 39.8 [24–59] mmHg; PVR 6.69 [2–15] WU | CYC (n=1); MMF+RTX (n=1); MMF+PDN (n=1); none (n=1) | Explanted lungs at time of transplant, end-stage SSc-ILD |
| Lung | Papazoglou et al. 2022 (GSE212109) | 3 (1 age/sex NR) | 52.0 [42–62] (n=2) | 66.7 (2/3) | SSc-ILD 100% | NR | NR | NR | NR | NR | NR | Demographic/technical metadata only; no clinical severity data available |
| Skin | Khanna et al. 2022, tofacitinib trial (GSE209635) | 15 (DB trial) | 50.8 ± 14.0 | 66.7 | dcSSc 100% | 2.1 ± 1.2 | 23.3 ± 8.4 | 88.4 ± 16.6 / 85.6 ± 19.8 | NR | Tendon friction rubs 13.3%; large joint contractures 33.3% | MMF 86.7%; MTX 13.3%; prednisone 26.7% | RCT-eligible; active, early diffuse cutaneous SSc |
| Skin | Tabib et al. 2021 (GSE138669) | 12 | NR (not reported) | 58.3 | dcSSc 100% | 2.42 [0.48–6.48] | 26.2 [12–43] | NR | NR | Histological fibrosis/inflammation/myofibroblast score (ND for most) | MYC/MTX/HCQ/D-Pen, variable per patient; 2/9 reported none | Biopsy-confirmed active skin disease |
| PBMC | Shimagami et al. 2025 (E-GEAD-872) | 21 | 64.9 [46–82] | 76.2 | dcSSc 14.3% / lcSSc 85.7% | 8.2 [0.3–32] | 7.0 [2–33] | NR | RNA pol III 42.9%; ACA 33.3%; ATA 14.3%; ANA 9.5% | SRC 19.0%; ILD 57.1%; PAH 14.3%; CIPO 9.5% | None, all treatment-naive | Broad clinical heterogeneity by design; treatment-naive at sampling |
| Synovium | Geiss et al. 2026, bioRxiv (unpublished) | 6 | 61.8 [55–68] | 50.0 | lcSSc 100% (localisation: wrist n=5, knee n=1) | 5.15 [0.15–21] | NR | NR | SCL-70 n=1; ACA n=2; RNA pol III n=2; anti-Ku+ANA n=1 | Pauci-immune 83.3%; lymphoid-myeloid 16.7% pathotype | cDMARD 50%; no therapy 50% | Active synovitis; |

**Legend / Abbreviations**: SSc, systemic sclerosis; CTRL, control; NR, not reported/not available in the source publication; ND, no data (assessed but not scored); dcSSc, diffuse cutaneous SSc; lcSSc, limited cutaneous SSc; Systemic sclerosis-associated interstitial lung disease (SSc-ILD); mRSS, modified Rodnan skin score; FVC, forced vital capacity; DLCO, diffusing capacity for carbon monoxide; mPAP, mean pulmonary arterial pressure; PVR, pulmonary vascular resistance; WU, Wood units; UIP, usual interstitial pneumonia; DB, double-blind; OLE, open-label extension; ACA, anti-centromere antibody; RNA pol III, anti-RNA polymerase III antibody; ATA, anti-topoisomerase I (Scl-70) antibody; ANA, anti-nuclear antibody; SRC, scleroderma renal crisis; ILD, interstitial lung disease; PAH, pulmonary arterial hypertension; CIPO, chronic intestinal pseudo-obstruction; CYC, cyclophosphamide; MMF, mycophenolate mofetil; RTX, rituximab; PDN, prednisone; MTX, methotrexate; HCQ, hydroxychloroquine; D-Pen, D-penicillamine; cDMARD, conventional disease-modifying antirheumatic drug; bDMARD, biological DMARD; NIC, non-inflammatory control (here: orthopedic trauma-derived synovial tissue); pathotype, histological synovial classification (pauci-immune vs. undefined/lymphoid); Krenn score, semi-quantitative histopathological synovitis score (range 0–9, combining lining layer hyperplasia, stromal cell density, and inflammatory infiltrate).

Supplementary figures

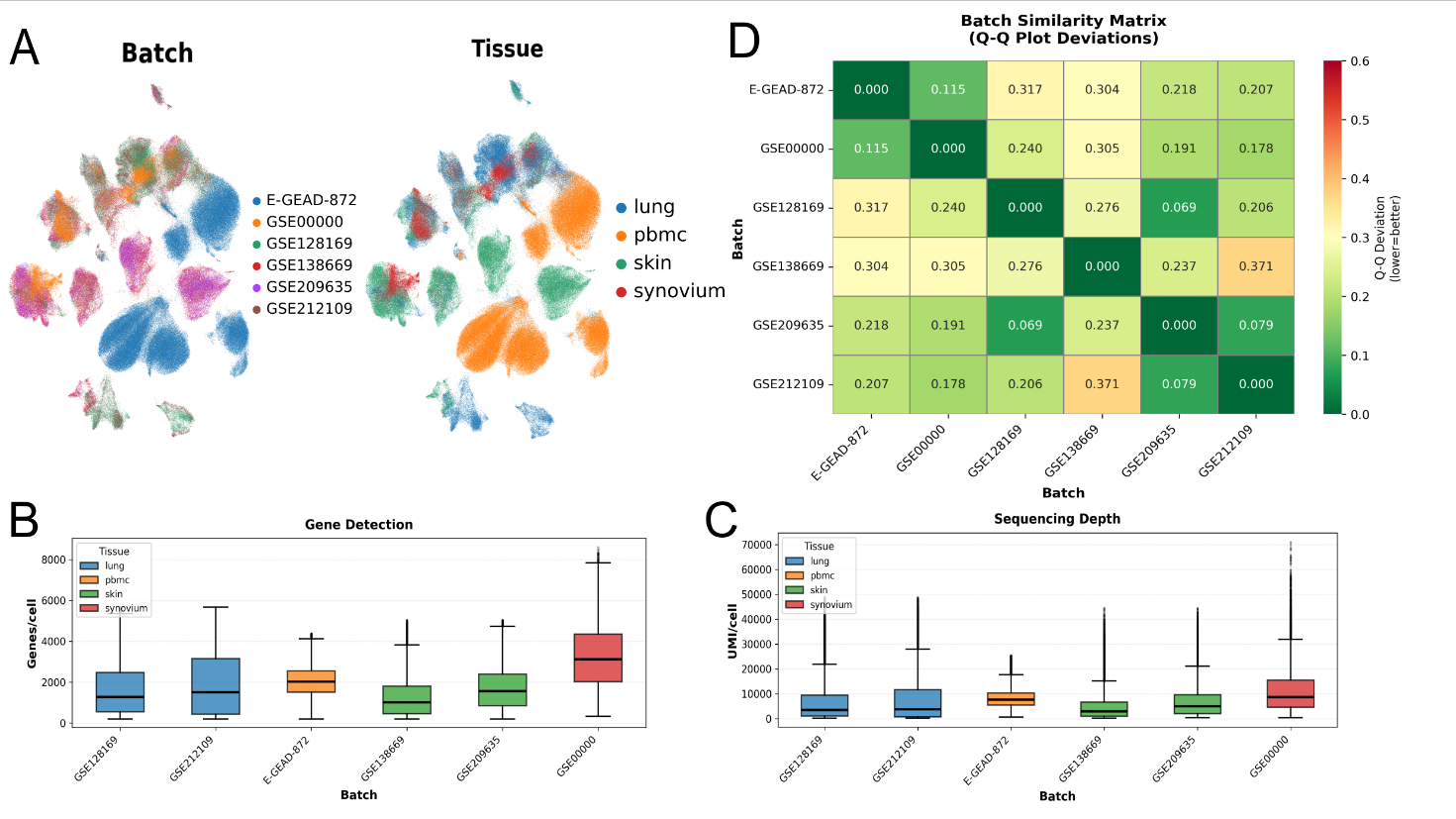

**Figure S1 Overview compatibility multi-organ single-cell transcriptomic atlas. A)** UMAP (Uniform Manifold Approximation and Projection) of the integrated atlas coloured by batch and sample source **B)** Boxplot showing atlas batches on the x-axis and the gene-per-cell ratio on the y-axis, with colours indicating the tissue of origin. **C)** Same as in (B), displaying the UMI-per-cell ratio on the y-axis. **D)** Batch similarity matrix based on Q–Q plot deviations, where matrix values represent the deviation across the expression value distribution for each pairwise batch comparison.

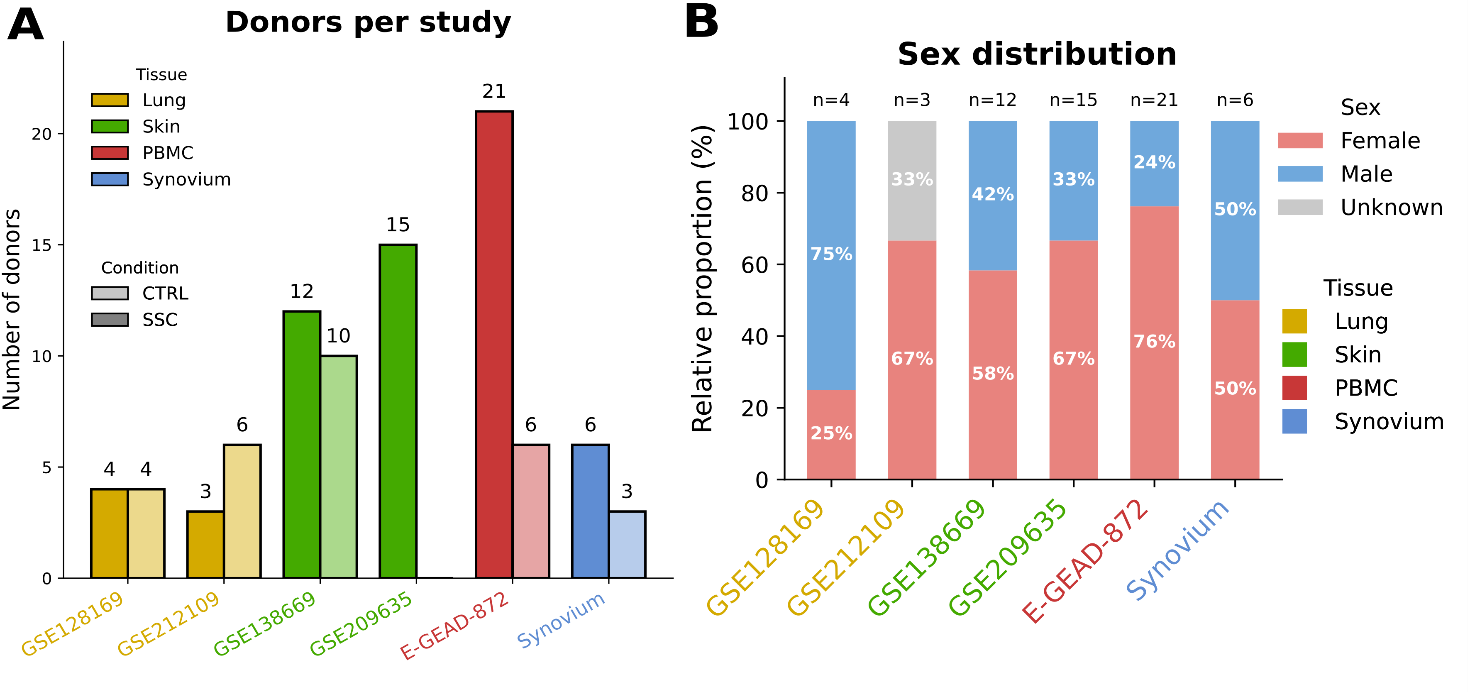

**Figure S2 Overview of the SSc multi-organ single-cell transcriptomic atlas. A)** Number of individuals per study stratified by disease condition (SSC, black outline; CTRL, grey outline). Bar fill colour indicates tissue of origin (yellow: lung; green: skin; red: PBMC; blue: synovium). **B)** Sex distribution per study (red: female; blue: male; grey: unavailable). Cohort name coloured by tissue of origin as in Figure A.

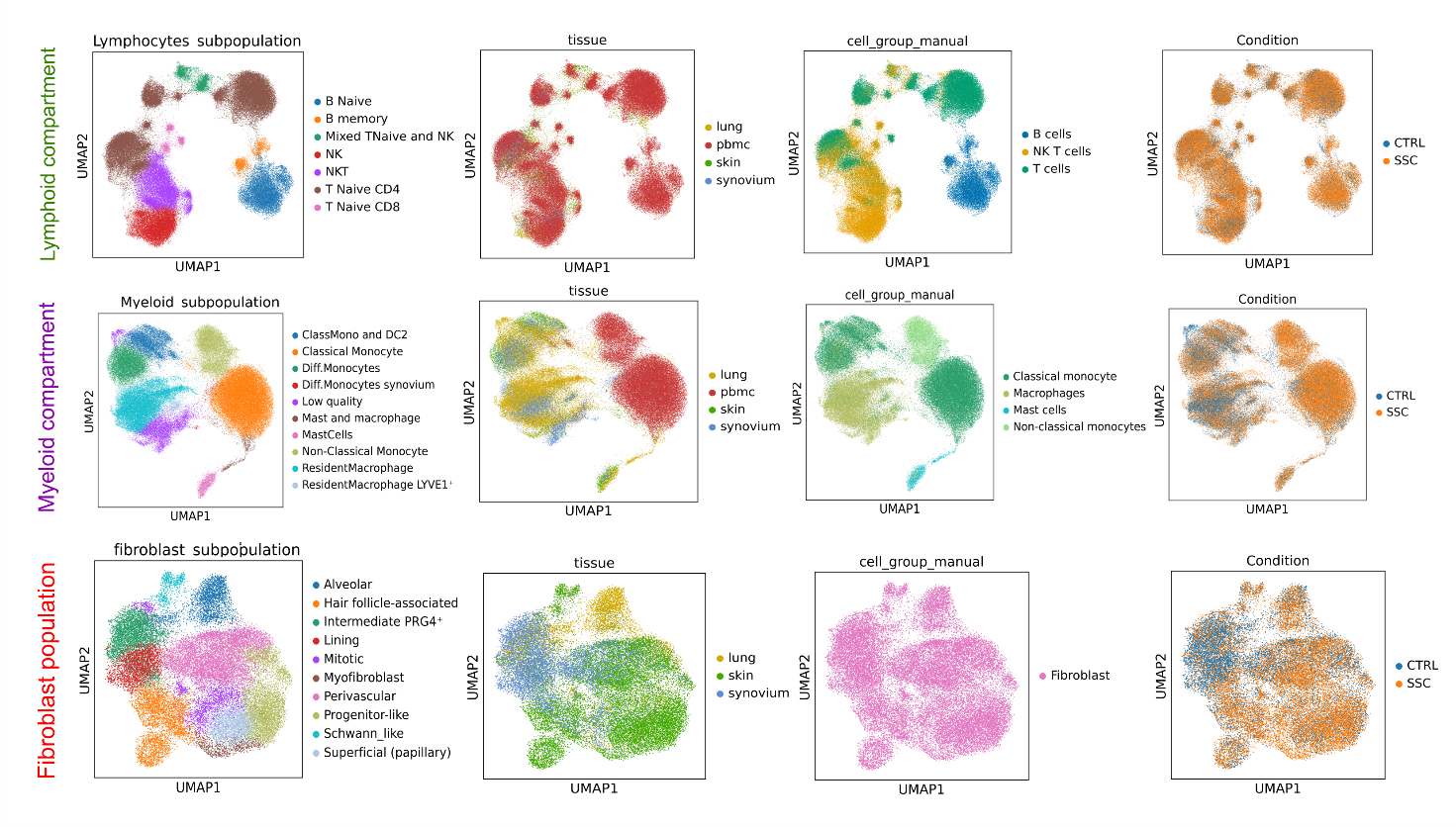
**Figure S3. UMAP visualization of independently reclustered lymphoid, myeloid compartments, and fibroblast population**. **A)** UMAP embeddings for the lymphoid (top row), myeloid (middle row) compartment, and fibroblast population (bottom row), reclustered independently of tissue of origin. For each compartment, panels from left to right show: subpopulation annotation; tissue of origin (lung, skin, synovium, PBMC); manually annotated cell type and disease condition (CTRL, SSC).
