## Supplementary Methods. for "Development of a multi-organ single-cell atlas of systemic sclerosis reveals shared and tissue-specific pathogenic pathways"

**Single-cell RNA sequencing datasets.**

To construct a multi-organ single-cell transcriptomic reference for systemic sclerosis (SSc), publicly available scRNA-seq datasets were retrieved from GEO and ArrayExpress. Lung data were obtained from two datasets (GSE128169 and GSE212109) [1,2], skin data from two datasets (GSE138669 and GSE209635) [3,4], and peripheral blood mononuclear cell (PBMC) data from one dataset (E-GEAD-872) [5]. Synovial tissue data were derived from our in-house cohort [6]. All datasets were processed independently prior to integration (**Table 1**).

**Quality control and preprocessing.**

Each publication dataset was preprocessed independently using Scanpy (v1.11) [7]. Quality control was performed by applying dynamic thresholds on the following per-cell metrics: mitochondrial RNA fraction (95^th^ percentile or < 20% if less than 20%), total UMI count (> 200), number of detected genes or UMI (from the 99^th^ percentile). Genes with less than 10 reads were excluded from the analysis. Doublets were identified and removed using Scrublet [8] and environmental RNA was corrected by Decontx package [9]. Two lung samples (SC281 and SC284) were excluded for low quality from the original study [2] as all the longitudinal samples in the datasets. In detail, were excluded the two E-GEAD-872 longitudinal samples of patient 1 (in the original study) from onset and after scleroderma renal crisis and the fifteen from GSE209635 after 6 weeks of Tofacitinib treatment. The 5 replicates of GSE128169 from different lung areas were included but merged as same individual for PyDESeq2, GSEA, gene set scores and their downstream analyses. Gene expression counts were normalised to 10,000 counts per cell followed by natural-log transformation, ln(1 + x).

**Assessment of cross-dataset comparability.**

Prior to integration, the comparability of normalised expression distributions across datasets was systematically evaluated. The mean and standard deviation of normalised log-transformed counts were computed for each dataset and compared through density plots and quantile–quantile (Q-Q) plots to assess distributional similarity (**Figure S1B-D**). Potential residual batch effects were further characterised by examining differences in per-cell gene count and UMI count distributions across samples. These analyses confirmed sufficient cross-dataset comparability as a prerequisite for multi-dataset integration. Two thousand highly variable genes were selected, and the number of genes and ribosomal and mitochondrial RNA content regressed out. Principal component analysis (PCA) was performed on the scaled data, followed by Harmony integration with default parameters [10]. The neighbourhood graph was constructed using the Harmony-corrected factors with n_neighbors = 15 and n_pcs = 50, followed by UMAP dimensionality reduction.

**Cell-type annotation.**

Cell-type annotation was performed using CellTypist (v1.7.1) [11], applying tissue-appropriate reference models to each tissue object. Specifically, the Human_IPF_Lung model was applied to lung datasets [12], the Adult_Human_Skin model to skin dataset [13], and the Immune_All_High model to the PBMCs dataset [14]. Synovium annotation was inherited by our previous study [6]. Automated annotations were subsequently reviewed and manually curated in the integrated object by marker genes to resolve ambiguous or inconsistent assignments and to harmonize nomenclature across tissue compartments. The final annotated integrated object served as the input for the interactive atlas application. To facilitate downstream analysis, cell types were also grouped into five cell compartments: Lymphoid (B and T lymphocytes, plasma cells, and NK T cells), Myeloid (classical and non-classical monocytes, mast cells, and DC2), Epithelial (differentiated and undifferentiated keratinocytes, multiciliated cells, AT1-AT2, and other epithelial cells), Mesenchymal (fibroblasts and pericytes), and Endothelial (endothelial cells). Schwann cells and melanocytes, present exclusively in the skin dataset, were excluded from downstream analyses.

**Interactive atlas application.**

The integrated and annotated single-cell object was deployed as an interactive web-based application built using the Streamlit framework (v1.52.1) and hosted on a dedicated virtual machine within the institutional computational infrastructure. The application allows users to apply single or combined filters on gene expression thresholds and on available metadata fields, including sample identifier, disease condition, SSc subtype, tissue of origin, and annotated cell compartments and types. All downstream analyses are performed dynamically on the resulting filtered subset of cells. The platform is organised into six analytical tabs, each dedicated to a specific module.

Tab 1 provides dataset-level exploration through UMAP visualization and distribution plots. Tab 2 enables gene-level expression analysis, including UMAP, violin plot, and dot plot representations for individual genes, with an option to compare expression between two user-defined groups. Tab 3 is dedicated to differential gene expression (DGE) analysis: pairwise comparisons are performed using the Wilcoxon rank-sum test or pseudobulk by counts sum. The results are visualised as interactive volcano plots, and pathway-level interpretation is supported through overrepresentation analysis via EnrichR [15] and Gene set enrichment analysis (GSEA) applied to MSigDB Hallmark, Reactome, and Gene Ontology Biological Process gene set collections [16]. In this manuscript only the pseudobulk method was applied alongside all the analyses. Tab 4 focuses on metadata-level associations, with Pearson or Spearman correlation for continuous variables and Wilcoxon-tested box plots for categorical comparisons, all performed on pseudobulk expression values computed as the mean gene expression per individual within each annotated cell type. Tab 5 provides a pathway-level analytical module using sc.tl.score_genes from pseudobulked samples, accepting up to three simultaneous metadata variables as grouping factors to enable stratified violin plots. Tab 6 serves as an administrative monitoring module displaying real-time computational resource usage.

**Pseudobulk differential gene expression and pathway enrichment analysis.**

To characterize transcriptional differences between SSc patients and CTRL at the level of individual cell populations, pseudobulk differential gene expression analysis was performed using PyDESeq2 v0.5.2 [17]. For each of the five annotated cell compartments in each tissue and for each cell type, gene expression counts were aggregated per individual to generate pseudobulk profiles. Differential expression was performed using the negative binomial model implemented in PyDESeq2, with disease condition as the primary contrast and low-count genes removed by independent filtering. GSEA was subsequently applied to the ranked gene list (ranked by the composite score derived from log2 fold change × −log10 adjusted p-value) using the Reactome gene set collection, with a false discovery rate (FDR) threshold of 0.05. Normalised enrichment scores (NES) > 0 indicate upregulation in SSc relative to CTRL; NES < 0 indicate downregulation. Significant pathways (FDR < 0.05) were subsequently annotated using the Reactome pathway hierarchy, mapping each pathway to its top-level (Level 1) and sub-level (Level 2) categories as defined in the curated Reactome ontology (reactome.org). For each of the most shared pathways, up to 5 genes were selected for display in **Figures 1G, 2D** and **3B**, ranked primarily by the number of organs (or cell groups/cell types) in which the gene was significantly shared, and secondarily by the number of significant pathways associated with that gene.

**Subpopulation reclustering of lymphoid, myeloid, and fibroblast compartments.**

To investigate the immune compartments and fibroblast population in greater detail, cells annotated as fibroblasts (fibroblast population); B cells, T cells, and NK T cells (lymphoid compartment); and classical monocytes, non-classical monocytes, mast cells, and macrophages (myeloid compartment) were extracted from the integrated multi-organ object and reclustered independently. Each subset was reprocessed using the same pipeline applied to construct the final integrated object, including regression of the number of genes and of mitochondrial and ribosomal RNA content, PCA, and Harmony integration. Clustering was performed using the Leiden algorithm, with a resolution of 0.4 for the lymphoid compartment, 0.8 for the myeloid compartment, and 1.5 for fibroblasts. Subpopulations presenting different subpopulation markers were excluded as doublets or mixed subclusters, the same for subclusters arise by cells with low counts. Tissue-specific subclusters were identified when at least 75% of cells within a given subcluster shared the same tissue of origin, as done in similar work [18]. To account for differences in the total number of cells captured per tissue within each compartment, we additionally required the normalised subcluster proportion, defined as the number of subcluster cells divided by the total number of cells of the same compartment captured in that tissue, to be at least twice as high in the dominant tissue as in each of the other tissues. Pseudobulk differential gene expression and GSEA were performed as described above for the main atlas, using PyDESeq2 and the Reactome gene set collection, comparing SSc to CTRL within each resulting subpopulation.

**Organ-level pseudobulk analysis**

The pseudobulk differential gene expression and GSEA were also performed on all cells aggregated per individual within each tissue, using the same PyDESeq2 and GSEA pipeline described above, without stratifying by cell group or cell type.

**Cross-resolution pathway score analysis**

A focused analysis on the cytokine, fibrotic pathways and OXPHOS signatures was performed. To this aim, gene set scores (sc.tl.score_genes, Scanpy) were computed for every cell within the lymphoid, myeloid, and fibroblast populations using the following Reactome gene sets. For fibrosis: collagen formation (R-HSA-1474290), TGF-beta signalling (Hallmark: M5896) and epithelial-mesenchymal transition (Hallmark: M5930) were considered; for the cytokine signature: interferon type I and II interferon signalling (Hallmark: M5911 and M5913), TNF signalling (Hallmark: M5890), IL-6 JAK-STAT3 signalling (Hallmark: M5897), IL-2 STAT5 signalling (Hallmark: M5947). Finally, for the OXPHOS score the Hallmark: M5936 (Oxidative phosphorylation) gene set was used. Scores were compared between SSc and CTRL at three levels of resolution: the compartment level (the five annotated cell compartments), the cell-type level (the manually curated cell types within each relevant cell group), and the subcluster level, using the same lymphoid, myeloid, and fibroblast subpopulation assignments described above. Endothelial and epithelial cell groups and cell types were additionally tested at the cell-group and cell-type level only, since a dedicated subclustering pipeline was not applied to these compartments. For each level, scores were aggregated to patient-level pseudobulk (mean score across a patient's cells within each group, restricted to patient-group combinations with ≥50 cells) to avoid pseudoreplication, and compared using the Mann-Whitney U test with Benjamini-Hochberg FDR correction. The 5 replicates of GSE128169 from different lung areas were merged as same individual as for the PyDESeq2, GSEA analysis.

References

1	Papazoglou A, Huang M, Bulik M, *et al.* Epigenetic Regulation of Profibrotic Macrophages in Systemic Sclerosis-Associated Interstitial Lung Disease. *Arthritis Rheumatol*. 2022;74:2003–14. doi: 10.1002/art.42286

2	Valenzi E, Bulik M, Tabib T, *et al.* Single-cell analysis reveals fibroblast heterogeneity and myofibroblasts in systemic sclerosis-associated interstitial lung disease. *Ann Rheum Dis*. 2019;78:1379–87. doi: 10.1136/annrheumdis-2018-214865

3	Khanna D, Padilla C, Tsoi LC, *et al.* Tofacitinib blocks IFN-regulated biomarker genes in skin fibroblasts and keratinocytes in a systemic sclerosis trial. *JCI Insight*. 2022;7:e159566. doi: 10.1172/jci.insight.159566

4	Tabib T, Huang M, Morse N, *et al.* Myofibroblast transcriptome indicates SFRP2hi fibroblast progenitors in systemic sclerosis skin. *Nat Commun*. 2021;12:4384. doi: 10.1038/s41467-021-24607-6

5	Shimagami H, Nishimura K, Matsushita H, *et al.* Single-cell analysis reveals immune cell abnormalities underlying the clinical heterogeneity of patients with systemic sclerosis. *Nat Commun*. 2025;16:4949. doi: 10.1038/s41467-025-60034-7

6	Geiss C, Cebrian CC, Houtman M, *et al.* Synovitis in systemic sclerosis is an interferon-driven stromal condition distinct from rheumatoid arthritis. 2026;2026.06.30.733140.

7	Wolf FA, Angerer P, Theis FJ. SCANPY: large-scale single-cell gene expression data analysis. *Genome Biol*. 2018;19:15. doi: 10.1186/s13059-017-1382-0

8	Wolock SL, Lopez R, Klein AM. Scrublet: Computational Identification of Cell Doublets in Single-Cell Transcriptomic Data. *Cell Systems*. 2019;8:281-291.e9. doi: 10.1016/j.cels.2018.11.005

9	Yang S, Corbett SE, Koga Y, *et al.* Decontamination of ambient RNA in single-cell RNA-seq with DecontX. *Genome Biol*. 2020;21:57. doi: 10.1186/s13059-020-1950-6

10	Korsunsky I, Millard N, Fan J, *et al.* Fast, sensitive and accurate integration of single-cell data with Harmony. *Nat Methods*. 2019;16:1289–96. doi: 10.1038/s41592-019-0619-0

11	Domínguez Conde C, Xu C, Jarvis LB, *et al.* Cross-tissue immune cell analysis reveals tissue-specific features in humans. *Science*. 2022;376:eabl5197. doi: 10.1126/science.abl5197

12	Adams TS, Schupp JC, Poli S, *et al.* Single-cell RNA-seq reveals ectopic and aberrant lung-resident cell populations in idiopathic pulmonary fibrosis. *Sci Adv*. 2020;6:eaba1983. doi: 10.1126/sciadv.aba1983

13	Reynolds G, Vegh P, Fletcher J, *et al.* Developmental cell programs are co-opted in inflammatory skin disease. *Science*. 2021;371:eaba6500. doi: 10.1126/science.aba6500

14	Domínguez Conde C, Xu C, Jarvis LB, *et al.* Cross-tissue immune cell analysis reveals tissue-specific features in humans. *Science*. 2022;376:eabl5197. doi: 10.1126/science.abl5197

15	Chen EY, Tan CM, Kou Y, *et al.* Enrichr: interactive and collaborative HTML5 gene list enrichment analysis tool. *BMC Bioinformatics*. 2013;14:128. doi: 10.1186/1471-2105-14-128

16	Subramanian A, Tamayo P, Mootha VK, *et al.* Gene set enrichment analysis: A knowledge-based approach for interpreting genome-wide expression profiles. *Proceedings of the National Academy of Sciences*. 2005;102:15545–50. doi: 10.1073/pnas.0506580102

17	Muzellec B, Teleńczuk M, Cabeli V, *et al.* PyDESeq2: a python package for bulk RNA-seq differential expression analysis. *Bioinformatics*. 2023;39:btad547. doi: 10.1093/bioinformatics/btad547

18	Domínguez Conde C, Xu C, Jarvis LB, *et al.* Cross-tissue immune cell analysis reveals tissue-specific features in humans. *Science*. 2022;376:eabl5197. doi: 10.1126/science.abl5197
